# Structural basis for GluA1 AMPA receptor regulation by PRRT1/SynDIG4 in LTP

**DOI:** 10.64898/2026.06.24.734257

**Authors:** Yaoyao Han, Xinyao Dou, Sew Peak-Chew, Sven Truckenbrodt, Josip Ivica, Ingo H. Greger

**Affiliations:** Neurobiology Division, Medical Research Council (MRC) Laboratory of Molecular Biology, Cambridge, UK; Protein C Nucleic Acids Chemistry (PNAC) Division, Medical Research Council (MRC) Laboratory of Molecular Biology, Cambridge, UK

**Author notes:** Correspondence: Ingo H Greger. These authors contributed equally (Yaoyao Han, Xinyao Dou).

## Abstract

AMPA receptor (AMPAR) signalling underlies long-term potentiation (LTP), a cellular mechanism for learning that selectively involves GluA1-type AMPARs through incompletely understood mechanisms. Here, we report a key role for the auxiliary subunit PRRT1/SynDIG4 in LTP. Cryo-EM structures of hippocampal AMPARs reveal preferential association of PRRT1 with GluA1 via its membrane-embedded CD225 domain, which sequesters the GluA1 C-terminus through a previously unexplored lipid modification on Cys825. Consequently, GluA1 regulatory motifs implicated in LTP, including the CaMKII binding site, are inaccessible. PRRT1 overexpression impairs LTP, which is rescued by the Cys825Ser mutation, presumably by restoring access of the GluA1 tail to Ser831 phosphorylation by CaMKII. Our data uncover a mechanism wherein PRRT1/SynDIG4 controls the availability of the GluA1 pool that is released upon the induction of LTP.

## Introduction

AMPA receptors (AMPARs) are the principal mediators of excitatory neurotransmission(*1*), their regulation is central to synapse potentiation and to learning and memory(*2–5*). AMPARs are multi-subunit complexes comprising a tetrameric core of the GluA1-4 subunits in various combinations, which is diversified further by auxiliary subunits that govern nearly all aspects of AMPAR operation, from channel gating to synaptic trafficking and anchoring at the post-synaptic density (PSD)(*1, 6*). While the structure and function of the canonical auxiliary subunits such as TARPs and CNIHs are increasingly well defined, the function and organisation of the ‘peripheral’ auxiliary subunits, including the Shisas and ‘di-spanins’ (PRRT1/SynDIG4 and PRRT2), are less well understood(*7, 8*).

NMDA receptor-dependent LTP is expressed through the activity-dependent insertion of GluA1-containing AMPARs into the synapse(*9–11*). Contrary to the predominantly synaptic GluA2/3 AMPARs, GluA1-containing receptors are supplied from extra-synaptic reserve pools outside the PSD upon LTP induction(*10, 12*), yet the mechanism governing this critical step remains unresolved. GluA1 uniquely accumulates at the cell surface driven by its sequence-diverse cytoplasmic C-terminus, which contains regulatory motifs and phosphorylation sites(*2, 13*). Phosphorylation of Ser845 by PKA increases the extrasynaptic pool of surface GluA1 in an activity-dependent manner(*2, 13*). During LTP induction, a key regulatory switch is phosphorylation of Ser831 by CaMKII(*14–17*), which is associated both with increased single-channel conductance and enhanced synaptic accumulation of GluA1(*18–20*), in part through direct interactions with activated CaMKII(*21*). In addition, phosphorylation of Ser816 and Ser 818(*2*) by protein kinase C (PKC) promotes binding of the scaffold protein 4.1N, thereby coupling AMPARs to the synaptic actin cytoskeleton(*22*). Lastly, synaptic anchoring depends on the sequence-diverse NTD within the synaptic cleft(*23, 24*).

PRRT1/SynDIG4 (hereafter PRRT1; proline-rich transmembrane protein 1) strongly localises with extra-synaptic GluA1and is implicated in regulating this GluA1 reserve pool(*25–27*). Here, we reveal the structural basis for selective PRRT1 association with GluA1 and show how this interaction sequesters GluA1 in a non-potentiated state. This mechanism involves a previously unrecognised lipid modification that ties the GluA1 C-tail to PRRT1 and restricts access of CaMKII, thereby blocking LTP. Synapse potentiation is restored upon de-lipidation of the GluA1 C-tail, which permits access to CaMKII.

## Results

### PRRT1/SynDIG4 associates with GluA1 in native AMPARs

LTP is most extensively studied at hippocampal synapses(*3, 5*). To study the AMPARs involved we purified the predominant GluA2-containing receptors from pig hippocampal synaptosomes (fig. S1A). Focussed processing of the sequence-diverse N-terminal domains (NTDs) in conjunction with an antibody fragment labelling the GluA1 NTD (11B8(*28*)) yielded three main receptor subtypes (Fig. 1A, fig. S1, B to D, and table S1): GluA1/2 di-heteromers (59%), GluA1/2/3 tri-heteromers (28%) and a smaller population of GluA2/3 di-heteromers (13%) (Fig. 1, A and D). These AMPARs have been associated with different synaptic trafficking and signalling functions(*10, 29*).

**Fig. 1.**
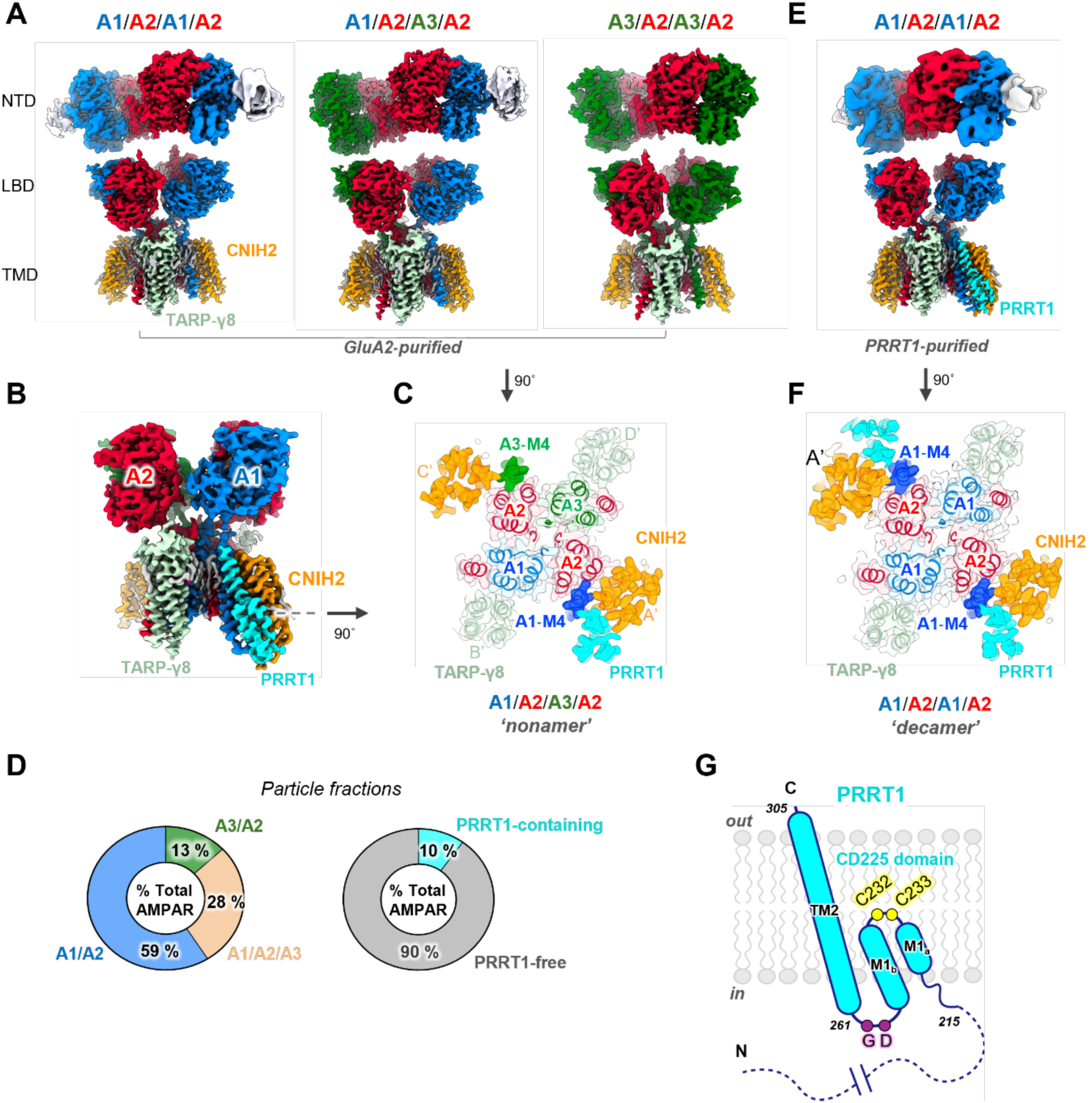
Main synaptic hippocampal AMPARs. **A,** Cryo-EM maps of native GluA2-containing AMPARs; subunit compositions were identified by NTD-masked and TMD-masked 3D classifications yielding three major classes: A1/A2/A1/A2, A1/A2/A3/A2, and A3/A2/A3/A2. Subunits are color-coded (A1, blue; A2, red; A3, green), with auxiliary proteins CNIH2 (orange) and TARP-γ8 (mint). Domain organization is indicated (NTD, LBD, TMD). **B,** Cryo-EM map of a PRRT1-bound AMPAR ‘nonameric’ complex. PRRT1 (cyan) occupies a single binding site between CNIH2 and GluA1 M4. **C,** Top-view slice through the TMD in panel A, illustrating the arrangement of core and auxiliary subunits within the receptor. Positions of the domain-swapped M4 helices are labeled. PRRT1 locates between GluA1 M4 and CNIH2. The A’, B’, C’, and D’ auxiliary subunit positions are denoted. **D,** Distribution of AMPAR subunit compositions (left) and fraction of PRRT1-containing particles (right). A1/A2 assemblies constitute the majority (59%), followed by A1/A2/A3 (28%) and A3/A2 (13%). Approximately 10% of total particles contain PRRT1. **E,** Cryo-EM map of a native PRRT1-containing AMPAR, purified with an anti-PRRT antibody. The ‘decameric’ receptor shows an A1/A2/A1/A2 receptor organization (A1, blue; A2, red) and lacks GluA3. **F,** Top-view slice through the TMD of the receptor decamer illustrating the spatial organization of the two PRRT1 subunits next to the GluA1 M4 helix and CNIH2. **G,** Schematic outline of PRRT1 topology. The transmembrane CD225-domain contains two transmembrane helices (M1a, M1b, and TM2) and a non-structured intracellular loop. Conserved residues and key structural features are indicated together with residue numbering. The disordered cytoplasmic N-terminus is indicated with a stippled line.

To reveal auxiliary subunit identity we included a ligand selectively marking TARP-γ8 in the preparation(*30, 31*) (fig. S1, E and F), a TARP heavily expressed across hippocampal synapse and essential for LTP expression(*3*). At nominal resolutions of ∼ 3 Å, we identified CNIH2 and TARP-γ8 at a 2:2 stoichiometry across the data set (Fig. 1C, fig. S2, A to E, and fig. S3A). Additional helical density adjacent to CNIH2 revealed the presence of the peripheral subunit PRRT1 (Fig. 1, B to D), which is further supported by mass-spectrometry (MS) (fig. S3, B to D). We observe one PRRT1 subunit per receptor, with its long C-terminal TM2 helix intersecting with CNIH2 and GluA1. Notably, PRRT1 is absent at the equivalent GluA3 position, suggesting selective binding to GluA1 (Fig. 1C, and fig. S3, B and C).

To clarify the apparent relationship between PRRT1 and GluA1, we used an anti-PRRT1 antibody for purification. The resulting AMPARs were now dominated by GluA1/2 di-heteromers associated with two PRRT1 subunits wedged between GluA1 and CNIH2 (Fig. 1, E and F, fig. S2, F and G, and fig. S4, A to D). These AMPAR ‘decamers’ further support PRRT1 selectivity for GluA1 over GluA3, and define the principles of AMPAR subunit organisation (movie S1): a tetrameric receptor core with two GluA2 subunits at the gating-dominant B/D positions flanked by two GluA1 subunits occupying the outer A/C sites. The ion channel core is encircled by two TARP-γ8 subunits at the B’/D’ sites and two CNIH2 subunits at the A’/C’ sites associated with two PRRT1 subunits (Fig. 1F and fig. S1F).

### The dimeric CNIH2-PRRT1 complex

PRRT1 is a Type-2 transmembrane protein composed of two domains. A cytoplasmic proline-rich N-terminus and a membrane-embedded C-terminal domain adopting a CD225 fold that comprises transmembrane helix 2 (TM2) and re-entrant helix 1 (M1a and 1b) (Fig. 1, B and G)(*32*). PRRT1 and PRRT2, another component of the AMPAR proteome (*33, 34*), belong to the ‘dispanin’ family of proteins implicated in membrane fusion events(*7, 32, 35, 36*).

PRRT1’s long TM2 helix adopts a (∼75°) tilt within the bilayer (Fig. 1, B and G) and is incompatible with a B’/C’ site location, where it would clash with TARP-γ8 (Fig. 2A). Instead, TM2 is embedded in a groove formed by GluA1 and CNIH2 where it extensively engages CNIH2 helices TM1 and TM4, as well as M4 of GluA1(Fig. 2, B and C). Key contact points include three residues unique to GluA1 (Fig. 2B, highlighted in pink). The CNIH2/PRRT1 dimer leaves an extensive ‘footprint’ on the receptor’s M1 and M4 helices (fig. S5A), which form the CNIH2 binding site and transmit gating modulation by CNIHs (Fig. 2C)(*37*). PRRT1 induces alterations in side chain geometry of CNIH2 Phe22 and Trp26 (Fig. 2D); these residues project into the interface between CNIH2 and the AMPAR and provide a possible mechanism underlying PRRT1-mediated gating modulation (see below).

**Fig. 2.**
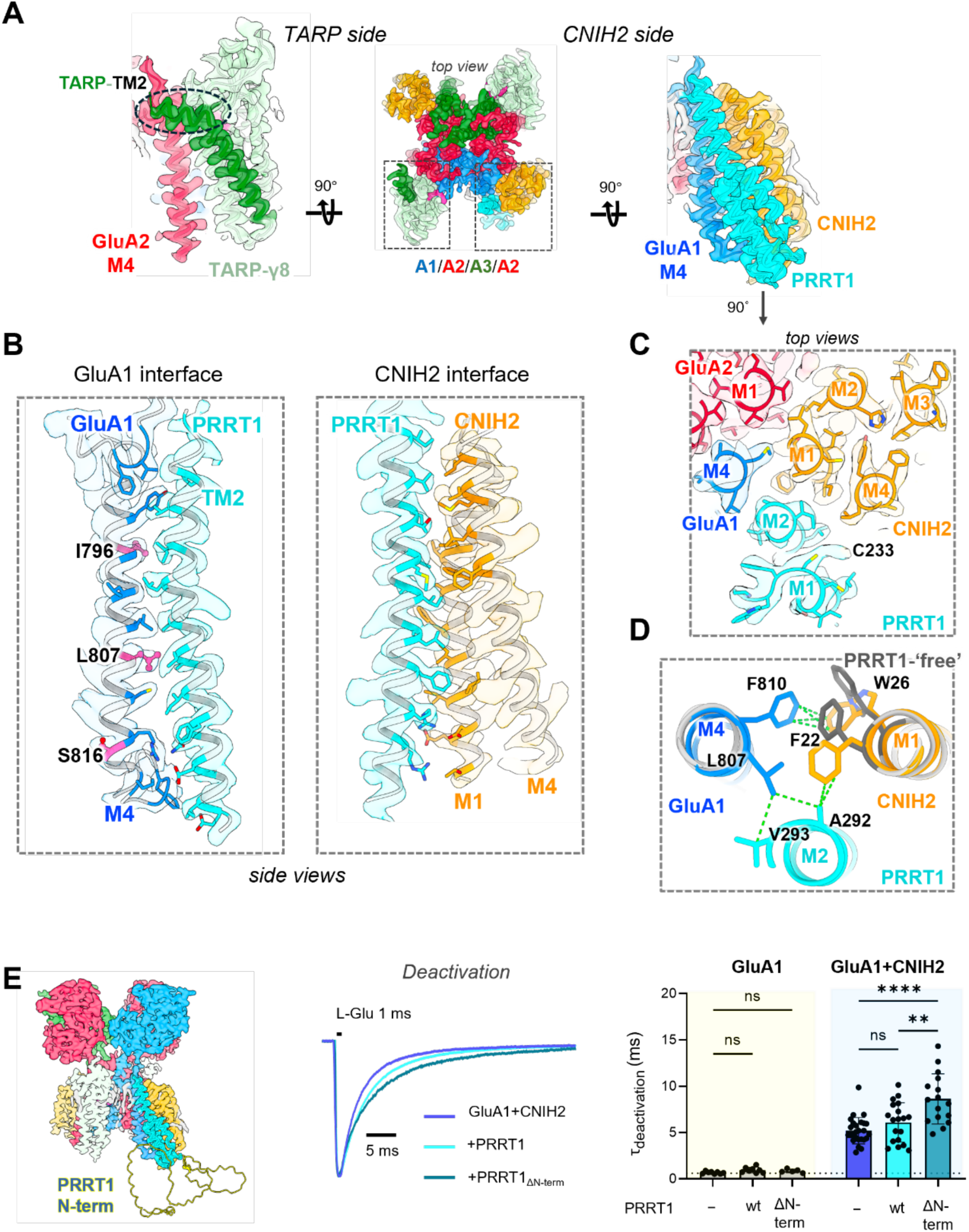
PRRT1-containing AMPAR organization and function. **A,** Center: top view onto a cryo-EM map of an A1/A2/A3/A2 receptor nonamer (color code as in Fig. 1). The TARP-γ8 (left) and CNIH2 sides (right) are boxed. Left panel: side view of the TARP-γ8 side, showing that the bend TM2 helix (black ellipsoid) would clash with PRRT1 TM2. Right panel: side view of the CNIH2 side harbouring PRRT1 next to GluA1 M4. **B,** Side views of interaction interfaces involving PRRT1. Left: between PRRT1-TM2 and the A1-M4 helix, highlighting three residues in pink (I796, L807, S816) unique to GluA1. Right: the PRRT1-M2 and CNIH2 interface involves the CNIH2 M1/M4 helices. **C,** Top view of the transmembrane arrangement showing the relative positions of PRRT1 helices (M1 and M2), A1 helices (M4), and CNIH2 helices (M1 and M4), illustrating the spatial organization of the interaction network. **D,** Close-up top views of residue-level interactions at the PRRT1–CNIH2–A1 interface. Comparison of PRRT1-bound (+PRRT1) and PRRT1-free A’/C’ sides shows repositioning of CNIH2 F22 and W26 by PRRT1. The hydrophobic interaction network involving A1 (F810, L807), CNIH2 (W26, F22), and PRRT1 (A292, V293) is depicted via green lines. **E,** Left: AMPAR A1/A2/A3/A2 nonamer including a PRRT1 AlphaFold model, depicting its disordered cytoplasmic N-terminus. Middle: representative current traces recorded from outside-out patches in response to 1 ms application of 10 mM glutamate from HEK293T cells expressing GluA1/CNIH2 or with PRRT1/ PRRT1_ΔN-term_, showing a slowing of current kinetics in the presence of PRRT1_ΔN-term_. Right: bar plots showing deactivation kinetics (1ms glutamate 10 mM) measured from outside-out patches excised from HEK293T cells expressing GluA1 alone or co-expressed with CNIH2, PRRT1, or PRRT1_ΔN-term_. Bar heights represent mean values. Top, entry into desensitization for GluA1 alone (n = 8 patches), GluA1+CNIH2 (n = 26 patches), GluA1+PRRT1 (n = 8 patches), GluA1+CNIH2+PRRT1 (n = 19 patches), GluA1+PRRT1_ΔN-term_ (n = 5 patches), and GluA1+CNIH2+PRRT1_ΔN-term_ (n = 16 patches). The effect of auxiliary subunits was assessed using One way ANOVA (F_(5, 63)_ = 32.90), P < 0.0001, followed by Šídák’s multiple comparisons test. **P = 0.0012; ****P<0.0001; ns, not significant (P > 0.05). Bar height represents the mean and the whickers SD. Dotted line represents the mean value of deactivation of GluA1.

The remainder of the PRRT1 CD225 domain is formed by the M1 helix, which is mostly polar, owing to this location at the membrane-cytosol boundary and splits into M1a and 1b (fig. S5B). The apex of M1 projects towards the center of the lipid bilayer and contains two conserved cysteine residues implicated in palmitoylation(*38*). A second highly conserved motif, Gly255/Asp256, resides within the cytosolic loop connecting M1b with TM2 (Fig. 1G) and has been linked to dispanin oligomerisation(*36*).

### PRRT1 gating function

We validated the subunit arrangement (Fig. 1) by patch-clamp electrophysiology of GluA1 expressed in HEK293 cells together with PRRT1 alone or with combinations of TARP-γ8 and CNIH2 (Fig. 2E). A subtle PRRT1-mediated slowing of both deactivation and desensitization was apparent only when it was transfected together with CNIH2 but not when expressed alone or with TARP-γ8 (fig. S5C, and table S2), as predicted from the structure (Fig. 2A). Gating modulation is mediated exclusively by the transmembrane CD225 domain. In fact, GluA1 kinetics slowed even further with a PRRT1 mutant lacking its N-terminus (PRRT1_ΔN-term_) (Fig. 2E). This can be explained by inefficient translation of the PRRT1 wild type (wt), resulting from its numerous N-terminal poly-proline stretches (fig. S5D)(*39*). PRRT1_ΔN-term_ expression levels were indeed greatly elevated, and the mutant efficiently accumulated at the cell surface (fig. S5, E and F), implying control of PRRT1 expression levels via its proline-rich N-terminus. The slowing of gating kinetics apparent with GluA1 homomers are not seen with GluA3 homomers and are largely blunted in GluA1/2 heteromers associated with both CNIH2 and TARP-γ8 (fig. S5G, and table S2). Therefore, PRRT1 modulation is GluA1-specific and may be minimal for receptor heteromers.

### PRRT1 sequesters the GluA1 C-terminus

The PRRT1-CNIH2 transmembrane sector protrudes into the cytosol, where it engages the membrane-proximal GluA1 C-tail (Fig. 3A). The impact of PRRT1 on GluA1 conformation is immediately apparent when comparing the PRRT1-occupied with the PRRT1-free site (Fig. 3, B and C). In the absence of PRRT1 density of the GluA1 M4 helix terminates at Ser816 due to flexibility of the C-tail. By contrast, with PRRT1 present, GluA1 M4 extends by > 10 residues into a cavity formed by PRRT1 and CNIH2 (Fig. 3C, bottom panel). This structural feature is apparent for both GluA1 subunits within the GluA1/2 decamer harbouring two PRRT1 subunits, and may therefore represent a PRRT1-specific signature element (fig. S6A).

**Fig. 3.**
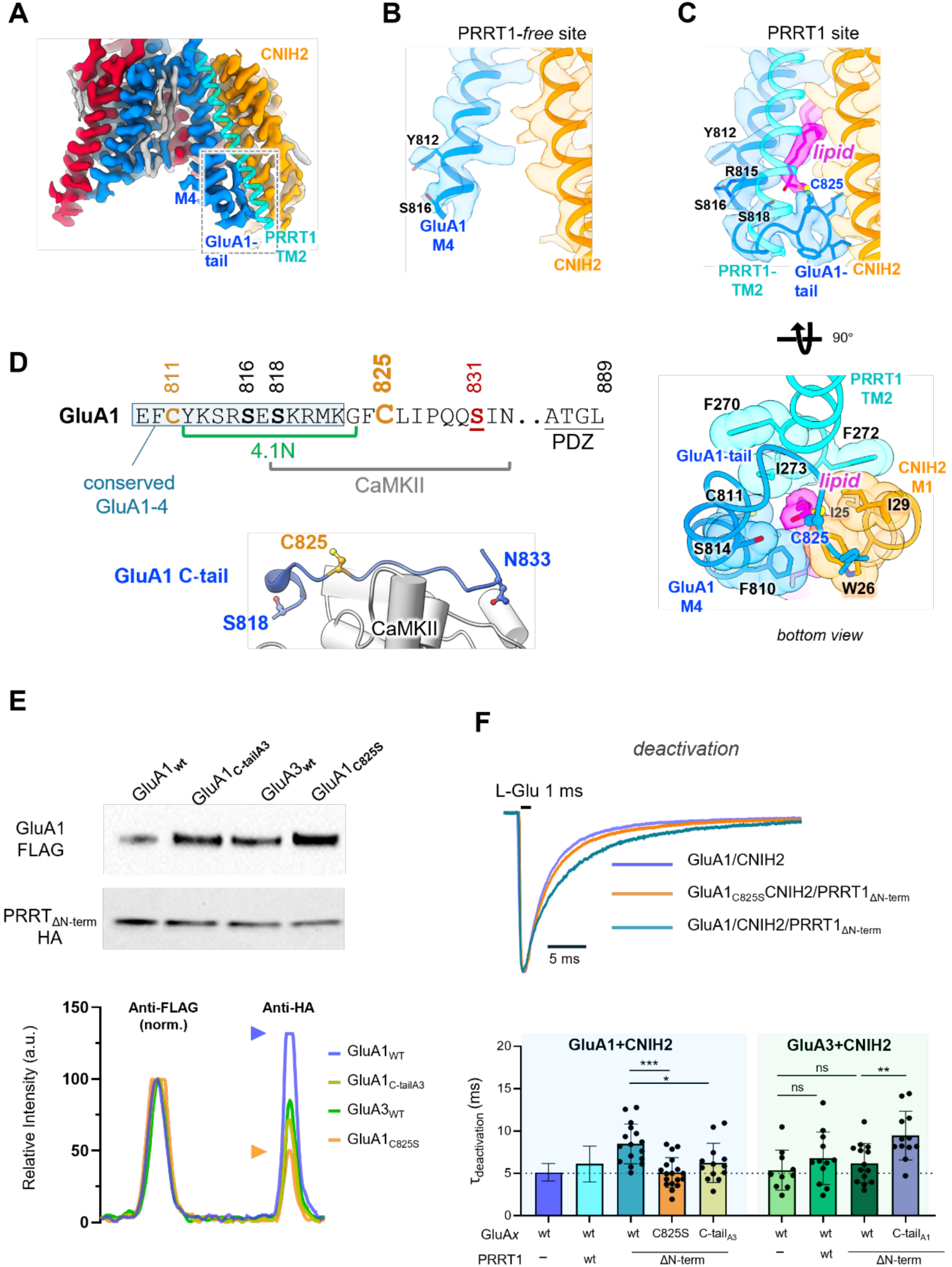
PRRT1 engages the GluA1 C-tail via a lipid. **A,** TMD cryo-EM density of a PRRT1-containing A1/A2/A1/A2 receptor nonamer. A model of the PRRT1 TM2 helix is shown in cyan. The boxed region indicates the extended density of GluA1-M4 tail shown further in panels b and c. **B-C,** Comparison between the PRRT1-free site (b) and the PRRT1-occupied site (c). In the absence of PRRT1, cryo-EM density for the GluA1 cytoplasmic tail terminates near Ser816. At the PRRT1 site, GluA1-M4 extends toward PRRT1 and CNIH2 and reveals a lipid-like density (pink) on GluA1 Cys825. Bottom view outlines the hydrophobic pocket formed by residues from GluA1-M4, PRRT1-M2, CNIH2-M1, and GluA2-M1 surrounding the lipid-like density (pink). **D,** Sequence and regulatory motifs within the GluA1 membrane-proximal C-tail. Residues implicated in post-translational modification and signaling are highlighted, including the 4.1N-binding motif, CaMKII interaction region, PKC phosphorylation sites Ser816/Ser818, CaMKII site Ser831, and palmitoylated cysteines Cys811 and Cys825 (described in this study). Residues conserved between GluA1-4 are boxed. Bottom: crystal structure of the CaMKII catalytic N-terminal domain (grey; PDB: 6X5Q) interacting with the GluA1 C-tail region Ser818 – Asn833 (blue). **E,** Interaction of PRRT1_ΔN-term_ with GluA1 and GluA3 C-terminal variants. Anti-FLAG immunoblot showing pull-down of FLAG-tagged GluA1 WT, GluA1 carrying the GluA3 C-terminal tail (GluA1_C-tailA3_), GluA3_WT_, and GluA1_C825S_. Co-immunoprecipitated HA-tagged PRRT1_ΔN-term_ was detected by anti-HA immunoblotting. Replacement of the GluA1 C-terminal tail with that of GluA3 or mutation of Cys825 reduced PRRT1_ΔN-term_ association compared with wild-type GluA1. The bottom panel represents quantification of band intensities (anti-FLAG signal was used for normalization, see Methods). The orange arrowhead depicts the reduction of GluA1_C825S_ interaction with PRRT1_ΔN-term_ compared to GluA1 wt (purple arrowhead). **F,** Top: Representative current traces recorded from outside-out patches in response to 1 ms application of 10 mM glutamate from HEK293T cells expressing GluA1/CNIH2, PRRT1/ PRRT1_ΔN-term_ and GluA1_C825S_/CNIH2 PRRT1_ΔN-term_, showing a slowing reduced effect of PRRT1_ΔN-term_ on C825S mutant. Bottom: bar plots showing deactivation kinetics (1ms glutamate 10 mM) measured from outside-out patches excised from HEK293T cells expressing GluA1 constructs (blue background) and GlluA3 (green background) co-expressed with CNIH2, PRRT1, or PRRT1_ΔN-term_. GluA1+CNIH2+PRRT1_ΔN-term_ (n = 16 patches), GluA1_C825S_+CNIH2+PRRT1_ΔN-term_ (n = 18 patches), GluA1_Ctail-A3_+CNIH2+PRRT1_ΔN-term_ (n = 13 patches), GluA3+CNIH2 (n = 10 patches), GluA3+CNIH2+PRRT1 (n = 12 patches), GluA3+CNIH2+PRRT1_ΔN-term_ (n = 14 patches) and GluA3_Ctail-A1_+CNIH2+PRRT1_ΔN-term_ (n = 12 patches). The effect of auxiliary subunits was assessed using One way ANOVA F_(7, 100)_ = 6.567, P < 0.0001, followed by Šídák’s multiple comparisons test. *P = 0.0494,**P = 0.0025; ***P = 0.0002, ns = not significant (P > 0.05). Bars and whiskers as in Fig. 2e.

### A lipid at GluA1 Cys825 engages PRRT1

We observe a lipid modification in the PRRT1-engaged GluA1 C-tail emanating from Cys825 (Fig. 3C). This residue has been proposed to be S-palmitoylated based on S-acyl proteomics(*40*), and O-nitrosylated(*41*). A C-16 palmitate fitted this density satisfactorily (fig. S6, B and C) and is further supported by a favourable GPS score(*42*), comparable to validated GluA1 palmitoylation sites(*43*). Hence, the GluA1 C-tail appears to harbour an additional palmitate downstream of the conserved Cys811 palmitoylation site(*43*) (Fig. 3D). This lipid inserts into a hydrophobic pocket formed by PRRT1-TM2, CNIH2-TM1 and GluA2-M1, anchoring the membrane-proximal GluA1 C-tail (fig. S6D).

Critically, Cys825 is surrounded by elements involved in regulating GluA1 synaptic function (Fig. 3D)(*2*). These include protein kinase (PKC) phosphorylation sites Ser816 and Ser818, which are embedded within a binding motif for protein 4.1N that anchors GluA1 to the synaptic actin cytoskeleton(*22, 44*). Further downstream lies Ser831, whose phosphorylation by CaMKII (and PKC) increases GluA1 single-channel conductance following NMDA receptor (NMDAR) activation(*2, 13*). PRRT1 will restrict access of 4.1N, PKC and CaMKII (Fig. 3C, and fig. S6C), thereby limiting phosphorylation and disrupting 4.1N-mediated anchoring as well as coupling to activated CaMKII, which forms a structural platform for GluA1 at potentiated synapses(*21, 45, 46*). Steric block of these motifs implicated in LTP offers a mechanism for PRRT1’s predicted role as an extra-synaptic ‘holding peg’ for GluA1-containing AMPARs(*7, 25–27*).

Further support for this mechanism is provided by immunoprecipitation of GluA1 from mouse brain. PRRT1 is efficiently co-precipitated with an antibody recognizing total GluA1 but not by an antibody specific for GluA1 phosphorylated at Ser831, with phosphorylation reporting unhindered access of CaMKII in the absence of PRRT (fig. S6E). Moreover, greater levels of Ser831-phosphorylated GluA1 are obtained from PRRT1 knock-out mouse brain(*47*).

### Cys825 determines PRRT1’s selectivity for GluA1

Cys825 is unique to GluA1 and absent in GluA2-4; its lipid modification may be the main contributor to PRRT1’s selective association with GluA1, in addition to Ile796 and Leu807 in the M4 helix (Fig. 2B). In support of this, both the Cys825Ser mutation and replacement of the GluA1 C-tail (from Arg815 onwards, Fig. 3D) with that of GluA3 reduced association with PRRT1 when immunoprecipitated from HEK293 cell extracts, to levels comparable to those observed with the GluA3-PRRT1 complex (Fig. 3E). The Cys825Ser mutation also blunted the kinetic component mediated by PRRT1_ΔN-term_, as did the GluA1 mutant harbouring the GluA3 C-tail. Conversely, transfer of the GluA1 tail onto GluA3 rendered the GluA3 chimera sensitive to modulation (Fig. 3F). Hence, the Cys825 lipid together with adjacent C-tail residues (Fig. 3C) contribute to the selective association of PRRT1 with GluA1.

### Extensive co-localisation of PRRT1 with GluA1 in hippocampus

To quantitatively assess the distribution of PRRT1 in neurons, we used expansion microscopy of ∼ 8-fold expanded mouse hippocampal slices combined with triple-colour immunostaining(*48*). First, we examined colocalization of PRRT1 with bona-fide excitatory synapses, visualised with pre-synaptic Bassoon and post-synaptic Shank 2. PRRT1 exhibited a wide distribution throughout the cell being detected in ∼ 70% of excitatory synapses, and within a large extra-synaptic pool encompassing ∼ 80% of total PRRT1 (fig. S7A), reflecting PRRT1 levels obtained by MS (fig. S3D). Next, we labelled PRRT1 and GluA1 alongside Shank 2 to determine the extent of PRRT1 co-localisation with GluA1 at synapses (Fig. 4A). The majority of the total GluA1-PRRT1 pool, ∼ 80%, is located extra-synaptic, lacking Shank 2 staining (Fig. 4B, left panel, and fig. S7B). This prominence of extra-synaptic PRRT1 has been reported(*25–27*) and is supported further by subcellular fractionation of pig brain (fig. S7C). Nevertheless, among GluA1-containing synapses, we detect a large proportion of PRRT1, ∼ 70%, highlighting its substantial presence at excitatory synapses, and its potential to regulate synaptic AMPARs.

**Fig. 4.**
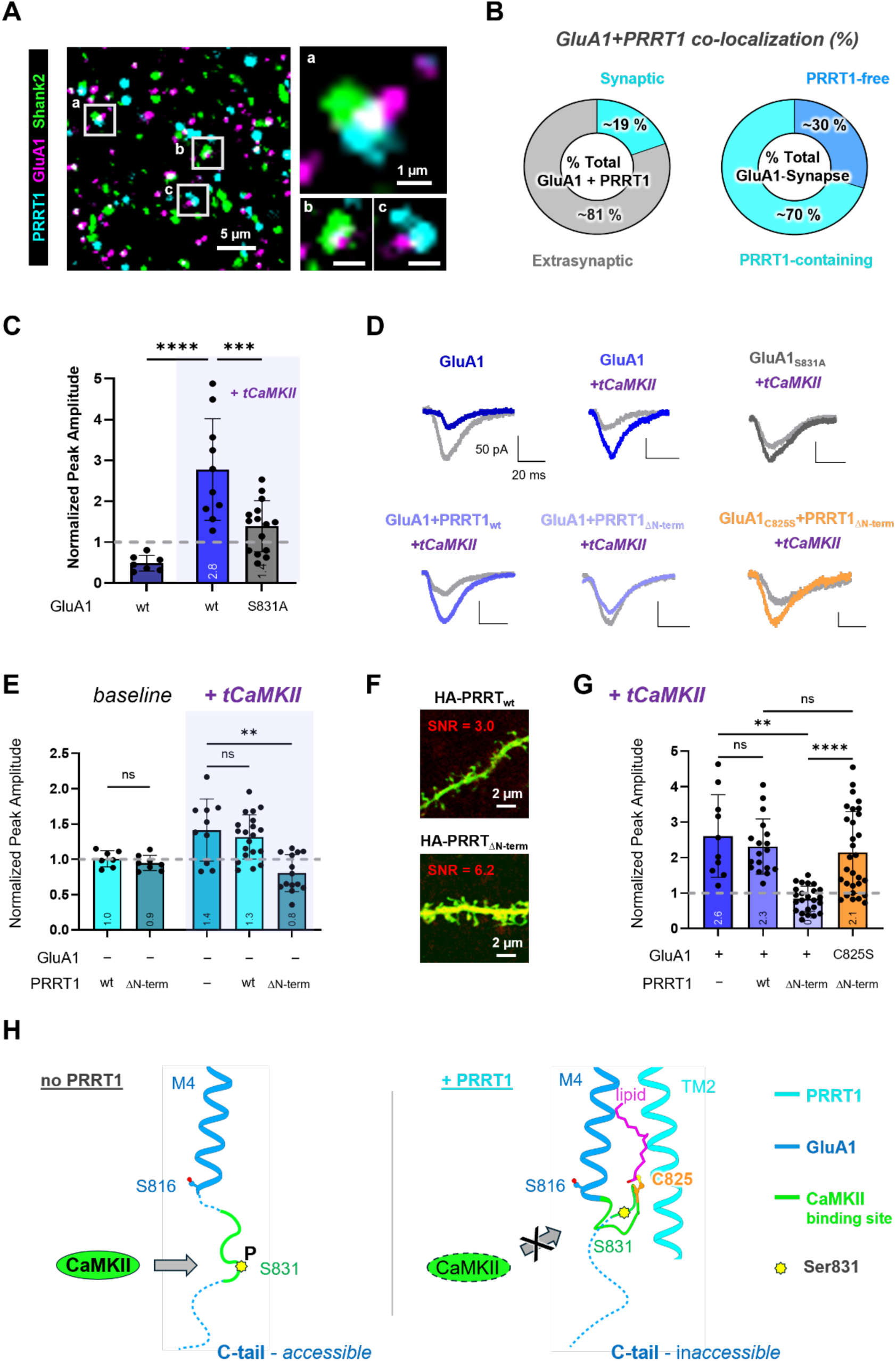
Synaptic distribution and function of PRRT1. **A,** Representative confocal images of ∼ 7-8-fold expanded mouse hippocampal slices staining with PRRT1 (cyan), GluA1 (magenta), and the postsynaptic marker Shank2 (green). Zoom-in panels show examples of a) synaptic colocalization of GluA1 and PRRT1 (cyan, magenta, and green), b) PRRT1-free GluA1-containing synapse (green and magenta), and c) extrasynaptic colocalization of GluA1 and PRRT1 (cyan and magenta). Scale bar for overview panel: 5 µm (biological scale: approximately 625 nm); Scale bar for zoom-in panels: 1 µm (biological scale: approximately 125 nm). **B,** Quantitative analysis of GluA1 and PRRT1 colocalization. Left: in the’ total GluA1 + PRRT1’ co-localised pool, synaptic and extra-synaptic portions are defined by the proximity with Shank2. Right: in the ‘total GluA1 synapse’ pool, PRRT1 presentence is defined by the proximity with PRRT1. Three biological replicates were performed (independent expansion and staining trials, each from different slices from different adult mice, n = 20 image stacks). **C,** Bar graph of normalized EPSC amplitudes from dual synaptic recordings in organotypic mouse hippocampal slice showing significance of GluA1_S831_ on truncated CaMKII (tCaMKII) potentiation (GluA1, n = 7; GluA1 with tCaMKII, n = 10; GluA1_S831A_ with tCaMKII, n = 15). Bars represent the mean, and error bars indicate SD. The effects of tCaMKII-induced potentiation on GluA1 wt and S831A mutant were assessed using Welch’s ANOVA (F_(2, 13.10)_ = 17.87, P = 0.0002), followed by Dunnett’s T3 multiple-comparisons test. *P < 0.05, **P < 0.01, ***P < 0.001, ****P < 0.0001; ns, not significant (P > 0.05). **D,** Representative EPSC traces of dual recordings from CA1 pyramidal neurons in organotypic hippocampal slices. Dual recordings were made on GluA1 alone, GluA1 / GluA1_S831A_ / GluA1+PRRT1_wt_ / GluA1+PRRT1_ΔN-term_ / GluA1_C825S_+PRRT1_ΔN-term_ with tCaMKII electroporated (blue and orange) cells with neighbouring untransfected (grey) cells. Scale bar: 50 pA, 20 ms. **E,** Bar graph showing normalized EPSC amplitudes from dual synaptic recordings in organotypic mouse hippocampal slice under baseline and truncated CaMKII (tCaMKII) potentiated condition (PRRT1_wt_, n = 7; PRRT1_ΔN-term_, n = 8; tCaMKII only, n = 10; PRRT1_wt_, n = 20; PRRT1_ΔN-term_, n = 14; GluA1, n = 20). Bars represent the mean, and error bars indicate SD. The effect of PRRT1 on native CA1 pyramidal neurons under basal or potentiated conditions was assessed using Welch’s ANOVA (F_(4, 24.06)_ = 10.52, P < 0.0001), followed by Dunnett’s T3 multiple-comparisons test. *P < 0.05, **P < 0.01, ***P < 0.001, ****P < 0.0001; ns, not significant (P > 0.05). **F,** Confocal images of transfected CA1 pyramidal neurons in organotypic slice. C-terminally HA-tagged PRRT1 wt and ΔN-term were electroporated with GFP as marker (green). Surface PRRT1 was visualized by immunostaining (red). Signal-to-noise ratio (SNR) was defined as foreground-to-background signal ratio (see Method). **G,** Bar graph showing normalized EPSC amplitudes from dual synaptic recordings in GluA1-overexpressed organotypic mouse hippocampal slice under tCaMKII potentiated condition (GluA1 alone, n = 10; GluA1+PRRT1_wt_, n = 18; GluA1+PRRT1_ΔN-term_, n = 24; GluA1_C825S_+PRRT1_ΔN-term_, n = 30). Bars represent the mean, and error bars indicate SD. The effect of PRRT1 on GluA1 co-expressed CA1 pyramidal neurons under potentiated condition was assessed using Welch’s ANOVA (F_(3, 34.31)_ = 14.07, P < 0.0001), followed by Dunnett’s T3 multiple-comparisons test. *P < 0.05, **P < 0.01, ***P < 0.001, ****P < 0.0001; ns, not significant (P > 0.05). **H,** Proposed molecular mechanism by which PRRT1 regulates CaMKII-mediated GluA1 Ser831 phosphorylation. Left panel: in the absence of PRRT1, the GluA1 C-tail remains accessible, enabling CaMKII binding and Ser831 phosphorylation. Right panel: in the presence of PRRT1, the GluA1 C-tail is anchored through interactions involving lipid-mediated Cys825 and the PRRT1 TM2 helix. This arrangement restricts access to CaMKII, reducing Ser831 phosphorylation and GluA1 synaptic anchoring. The CaMKII binding region on the GluA1 C-tail is indicated in green.

### PRRT1 impairs LTP via Cys825 lipidation

The extensive overlap of GluA1 with PRRT1 supports a functional role for this auxiliary subunit. Ser831 phosphorylation is widely associated with synaptic potentiation(*2, 13, 15, 17, 48, 49*) and PRRT1-mediated shielding of the CaMKII binding site (Fig. 3, C and D) could interfere with this process. To establish the functional impact of Ser831, we electroporated GluA1 wt and the GluA1_Ser831Ala_ phospho-null mutant together with constitutively active truncated CaMKII (tCaMKII) into organotypic hippocampal slices (fig. S7D). tCaMKII mimics and occludes LTP and has been used to probe the mechanisms underlying synaptic potentiation(*9, 50, 51*). In line with previous studies, tCaMKII strongly potentiated excitatory post-synaptic currents (EPSCs) in neurons expressing GluA1 wt, whereas this effect was attenuated in cells expressing GluA1_Ser831Ala_ (Fig. 4, C and D). Simultaneous recordings from untransfected neurons enabled normalisation and a direct comparison across conditions (Fig. 4C, broken horizontal line). Hence, Ser831 phosphorylation chiefly contributes to synapse potentiation by tCaMKII.

To assess PRRT1’s synaptic function we electroporated PRRT1 wt and PRRT1_ΔN-term_, the mutant expressing increased levels of the CD225 domain (fig. S5E). Baseline EPSCs were comparable with either of the two PRRT1 isoforms (Fig. 4E and fig. S7E). However, under potentiating conditions in the presence of tCaMKII, PRRT1_ΔN-term_ substantially reduced potentiation (Fig. 4E). This can be attributed to exogenous PRRT wt not substantially exceeding endogenous PRRT1 levels. By contrast, the elevated expression of PRRT1_ΔN-term_ unmasks the regulatory contribution of the CD225 transmembrane domain, with the mutant strongly accumulating at CA1 synapses (Fig. 4F). Indeed, when boosting EPSCs by co-expressing GluA1 with tCaMKII, we obtain only a trend-wise reduction of current amplitudes with PRRT wt, whereas PRRT1_ΔN-term_ caused severe reduction (GluA1 + PRRT1_ΔN-term_ vs. untransfected: 0.84 ± 0.36, n = 24), comparable to GluA1_Ser831Ala_ (Fig. 4G). Critically, the palmitoylation-deficient GluA1_Cys825Ser_ mutant almost completely restored potentiation in the presence of PRRT1_ΔN-term_ (Fig. 4G). Hence, PRRT1 engagement of essential GluA1 C-tail motifs attenuates synaptic potentiation and illuminates PRRT1’s role in regulating GluA1-type AMPARs in LTP (Fig. 4H).

## Conclusion

We present a mechanism by which the auxiliary subunit PRRT1 regulates signalling of GluA1 AMPARs (Fig. 4H). The preferential association of PRRT1 with the GluA1 subunit, revealed by native AMPAR structures, is mediated by the sequence-diverse GluA1 C-tail via lipidation of Cys825. Beyond recruiting PRRT1, this previously unrecognised modification sterically restricts key regulatory motifs within the GluA1 tail. Thus, the GluA1 C-tail recruits PRRT1 to the receptor, which determines access to motifs that govern GluA1 conductance and synaptic anchoring(*2, 13, 45*).

PRRT1 is a peripheral auxiliary subunit that, unlike TARPs, GSG1l and CNIHs, does not directly modulate AMPAR gating, consistent with its distinct binding mode. Instead of engaging the gate-surrounding M1 and M4 helices(*37*), PRRT1 contacts one face of GluA1 M4 and the membrane proximal GluA1 C-tail. Its modulatory activity is therefore indirect and mediated by CNIH2, whose structure is influenced by PRRT1. Notably, PRRT1 is the first auxiliary subunit to exhibit core subunit-selective activity, revealing emerging principles of AMPAR organisation (movie S1).

We propose a dual role for the PRRT1transmembrane CD225 domain: CNIH2-dependent gating modulation and AMPAR retention at extrasynaptic sites through masking of the GluA1-tail. However, the function of the disordered PRRT1 N-terminus remains unresolved. In the related PRRT2, the N-terminus has been shown to interact with the presynaptic SNARE machinery that regulates synaptic vesicle fusion(*32, 36*). In PRRT1, the N-terminus may facilitate sequestration of PRRT1-AMPAR complexes outside the PSD by engaging target sites that remain to be identified. Activity-dependent release of GluA1 from PRRT1 could be mediated by reversible palmitoylation of GluA1 Cys825, analogous to the dynamic regulation of PSD-95 and A-kinase anchoring protein (AKAP)79/150(*52–54*). Accordingly, Cys825 de-lipidation destabilises PRRT1 association with GluA1 and enable access of protein 4.1N, PKC and CaMKII to permit synaptic anchoring and potentiated transmission.

## Acknowledgements

We thank Nick Barry and Jerome Boulanger for support with imaging and data analysis. We thank Tom Smith for the analysis of the MS data. Paul Emsley, Lucrezia Catapano for helpful suggestions with model building and Coot figure presentation. We also thank Edward Ziff, Ondrej Cais and Trevor Rutherford for comments on the manuscript. We are grateful to the Biological Services teams at the LMB and Ares facilities, the LMB scientific computing, mass-spectrometry, light microscopy and the EM facilities for support.

## Data, code and materials availability

Cryo-EM density maps and corresponding atomic coordinates generated in this study have been deposited in the Electron Microscopy Data Bank (EMDB) and the Protein Data Bank (PDB), respectively, under the following accession codes: native GluA2 AMPAR A1A2A1A2 NTD, EMD-58413/PDB 31HF; native GluA2 AMPAR A1A2A3A2 NTD, EMD-58414/PDB 31HG; native GluA2 AMPAR A3A2A3A2 NTD, EMD-58415/PDB 31HH; native GluA2-containing AMPAR TMD octamer, EMD-58416/PDB 31HI; PRRT1-containing AMPAR nonamer TMD, EMD-58417/PDB 31HJ; PRRT1-containing AMPAR decamer TMD, EMD-58418/PDB 31HK; and PRRT1-containing AMPAR decamer NTD, EMD-58419. Custom code for 3D multiple spot colocalization is available at https://raw.githubusercontent.com/jboulanger/imagej-macro/refs/heads/main/Colocalization_Analysis/MultiChannel_Spot_3D_Colocalization.ijm.

Source data are provided with this paper.

## Author contributions

All authors conceived the study, supervised by IHG, who wrote the paper with input from the authors. YH performed protein purification and cryo-EM data collection, EM data processing and model building; XD performed and analysed synaptic recordings; JI performed and analysed HEK293 cell electrophysiological experiments. XD performed Expansion microscopy and imaging supervised by ST; SPC performed mass spectrometry and data analysis.

