## Supplementary data for "Structural basis for GluA1 AMPA receptor regulation by PRRT1/SynDIG4 in LTP"

**The PDF file includes:**

Materials and Methods
Figs. S1 to S7
Tables S1 to S2

**Other Supplementary Materials for this manuscript include the following:**

Movie S1
Data S1

#### Methods

##### Production and purification of 11B8

The open reading frame of 11B8 was synthesized (Genewiz) and cloned into the phLSEC vector with a C-terminal His tag(*28*). For transient expression, 2 mg of plasmid DNA was mixed with 6 mg of polyethylenimine (PEI), incubated briefly, and used to transfect suspension cells. Cells were cultured for 60–72 h post-transfection to allow secretion of the antibody into the culture medium. The conditioned medium was harvested and clarified by filtration through a 0.22 μm membrane.

The supernatant was buffer-exchanged into 20 mM Tris-HCl (pH 8.0), 150 mM NaCl using an ÄKTA Flux system and subsequently purified by Ni-NTA affinity chromatography. The antibody-containing solution was incubated with Ni-NTA resin, followed by washing with 20 mM Tris-HCl (pH 8.0), 150 mM NaCl, and 40 mM imidazole. Bound protein was eluted with 20 mM Tris-HCl (pH 8.0), 150 mM NaCl, and 250 mM imidazole.

Eluted fractions were concentrated using a 10 kDa molecular weight cut-off filter and further purified by size-exclusion chromatography on a Superose 6 Increase 10/300 GL column equilibrated in 20 mM Tris-HCl (pH 8.0), 150 mM NaCl. Peak fractions corresponding to 11B8 were collected for downstream applications.

##### Antibody production and purification

For expression and purification of antibodies, plasmids encoding Anti-GluA2 [L21/32R] antibody or Anti-SynDIG4/Prrt1 [L102/45R] antibody (P1316-IgG2a backbone engineered with a C-terminal twin-Strep tag)(*55*) were transiently transfected into Expi293F cells. Briefly, 2 mg of plasmid DNA was mixed with 6 mg of polyethylenimine (PEI) and incubated before transfection. Cells were cultured for 60–72 h post-transfection to allow secretion of antibodies into the culture medium.

The conditioned medium was harvested and clarified by filtration through a 0.22 μm membrane. Buffer exchange into 20 mM Tris-HCl, pH 8.0, 150 mM NaCl was performed using an ÄKTA Flux system. The antibody-containing solution was incubated overnight at 4°C with 1 mL of Strep-Tactin 4× Flow beads. After incubation, the beads were washed with

50 mL of wash buffer (20 mM Tris-HCl, pH 8.0, 150 mM NaCl). Bound antibodies were eluted with elution buffer (20 mM Tris-HCl, pH 8.0, 150 mM NaCl).
Eluted fractions were further purified by size-exclusion chromatography on a Superose 6 Increase 10/300 GL column equilibrated in 20 mM Tris-HCl, pH 8.0, 150 mM NaCl. Peak fractions corresponding to monomeric antibody were collected for downstream applications.

**Purification of native GluA2 AMPAR complexes**

AMPAR complexes were purified natively from freshly dissected pig hippocampi (animals >5 months old) following a procedure described recently(*56*). Brains were obtained from a local butcher and transported in ice-cold phosphate-buffered saline (PBS). Hippocampal tissues from six pig brains were dissected, pooled (23 g total), and homogenized in ice-cold homogenization buffer (320 mM sucrose, 4 mM HEPES, pH 7.5, 100 μM PMSF) using a blender (3–5 cycles of 5 s), followed by further homogenization with a Potter–Elvehjem homogenizer (10 strokes) until complete tissue disruption. The homogenate was diluted to 200 mL with homogenization buffer and centrifuged at 1,000 × g for 10 min at 4°C. The supernatant (S1) was collected and centrifuged at 12,000 × g for 35 min at 4°C (Beckman Ti-45). The resulting synaptosome pellet (P2) was resuspended in homogenization buffer, aliquoted, snap-frozen in liquid nitrogen, and stored at −70°C.
For protein purification, P2 fractions were thawed and hypotonically lysed in ice-cold water (1:9, v/v) for 15 min at 4°C under gentle rotation. The lysate was centrifuged at 25,000 × g for 30 min at 4°C (Beckman Ti-45), and the resulting pellet was solubilized in lysis buffer (150 mM NaCl, 20 mM Tris-HCl pH 8.0, 1% digitonin, 5 μM NBQX, 40 μM TARPγ8 ligand AMPA-IN-3 (MedChemExpress Cat# HY-175443), 100 μM PMSF, and protease inhibitors) for 3 h at 4°C. Insoluble material was removed by ultracentrifugation at 150,000 × g for 45 min at 4°C (Beckman Ti-45), and the supernatant was collected on ice.
For affinity purification, 1.0 mg of neuromab antibody (engineered with a twin-Strep tag) was added to the lysate and incubated for 4 h at 4°C under gentle rotation. Subsequently, 1 mL of Strep-Tactin XT 4Flow bead slurry was added and incubated overnight (~14 h) at 4°C.

Beads were washed with 30 mL wash buffer (150 mM NaCl, 20 mM Tris-HCl, pH 8.0, 0.01% GDN) using a gravity column. Bound complexes were eluted in eight fractions (0.5 mL each) using elution buffer (150 mM NaCl, 100 mM Tris-HCl, pH 8.0, 0.01% GDN, 50 mM biotin). Eluted fractions were concentrated using a 100 kDa molecular weight cut-off concentrator. For GluA2-containing AMPAR complexes, purified samples were incubated with 11B8 Fab (A1 probe) at 4°C for 2 h and further analyzed by size-exclusion chromatography on a Superose 6 Increase 10/300 GL column equilibrated in SEC buffer (150 mM NaCl, 20 mM Tris-HCl pH 8.0, 0.01% GDN). Peak fractions were collected, concentrated, and supplemented with 300 μM NBQX and 40 μM TARPγ8 ligand before cryo-EM grid preparation.

**Purification of native PRRT1 AMPAR complexes**

For purification using the PRRT1-NTD antibody, the procedure was performed as described above for the GluA2-CTD antibody. Due to low protein yield, size-exclusion chromatography was omitted, and eluted fractions were directly concentrated and used for cryo-EM grid preparation.

**Cryo-EM grid preparation and data collection**

Before vitrification, purified AMPAR complexes were incubated with 200 μM NBQX and 40 μM TARPγ8 ligand. for at least 30 min on ice. Ultrafoil Au 300 1.2/1.3 grids were glow-discharged before sample application. Subsequently, 3–4 μL of sample was applied to the grids and incubated for ~5 s, followed by blotting for 2 s with a blot force of −3. Grids were rapidly plunge-frozen in liquid ethane using a Vitrobot Mark IV (Thermo Fisher Scientific) operated at 4°C and 100% humidity.

Cryo-EM data were collected on a 300 kV Titan Krios G4 microscope (Thermo Fisher Scientific) equipped with a BioContinuum K3 Direct Electron Detector and a Gatan GIF energy filter (20 eV slit width). Data acquisition was performed using EPU with beam-image shift (AFIS enabled). Data collection was performed at a magnification of 105 kx in counted super-resolution mode with 2× binning, resulting in a pixel size of 0.826 Å per pixel. Movies were recorded for 40 frames and 1.8 s, resulting in a total dose of 40 e−/Å2. The defocus values ranged from −1.2 to −2.4 μm.

**Cryo-EM data processing and model building**

All cryo-EM data processing was carried out in cryoSPARC v4.7.1. For the native GluA2 AMPAR dataset, a total of ~32,000 movie stacks were subjected to Patch Motion Correction, followed by Patch CTF Estimation. Micrographs with poor CTF fits or excessive drift were discarded. Particle picking was performed using Template Picker, and particles were extracted with a box size of 512 pixels (Fourier cropped, bin 4), yielding 1.84 million particles.

Extracted particles were subjected to Heterogeneous Refinement to remove junk classes and separate distinct conformations, resulting in multiple classes. The best-resolved class (760k particles) was selected and re-extracted at full pixel size (box size 512) for further refinement. This particle set was refined using Non-uniform Refinement, producing an initial consensus reconstruction.

To resolve structural heterogeneity, particles were subjected to 3D Classification (heterogeneous refinement with multiple initial models) without masking, yielding several classes with distinct N-terminal domain (NTD) arrangements. Classes were grouped based on subunit composition (A1/A2/A3 distribution) and pooled accordingly. Selected particle subsets were further refined using Local Refinement with masks applied to specific regions (NTD or transmembrane domain, TMD).

For high-resolution reconstruction of domain-specific features, focused classification and refinement strategies were employed. For the NTD, particles were subjected to iterative Heterogeneous Refinement with an NTD mask, followed by Local Refinement, resulting in maps at 3.1 Å resolution. For the TMD region, particles were refined using a TMD-focused mask, yielding a map at 2.7 Å resolution. Additional rounds of 3D Classification further improved homogeneity, and the best-resolved class (65k particles) was refined to ~3.1 Å resolution.

Distinct conformational classes (Classes 0–7) were further grouped and refined independently. Selected classes (Classes 0/1/3/6/7, 2/4, and 5) were subjected to Local Refinement with masks applied to NTD or LBD/TMD regions, producing final reconstructions at resolutions ranging from 3.1 Å to 3.6 Å.

For the native PRRT1 AMPAR dataset, all data processing was performed in cryoSPARC. A total of 31,000 movie stacks were subjected to Patch Motion Correction and Patch CTF Estimation, followed by curation to remove low-quality micrographs. Particles were picked using Template Picker, extracted with a box size of 512 pixels (Fourier cropped, bin 4), and subjected to 2D Classification, yielding ~3.7 million particles, of which ~200,000 particles showing clear secondary structural features were selected for downstream analysis.

Selected particles were subjected to iterative Heterogeneous Refinement to remove junk and separate structurally distinct classes. The best-resolved class (~770k particles) was further refined through additional rounds of Heterogeneous Refinement, yielding a subset of 350k particles with improved structural features. This particle set was further cleaned through additional classification, resulting in 240k particles that were used for ab initio reconstruction to generate initial models.

Subsequent rounds of Ab-initio Reconstruction and classification identified a dominant class (76k particles) corresponding to intact PRRT1–AMPAR complexes. These particles were re-extracted at full pixel size (box size 512) and subjected to a second round of Ab-initio Reconstruction, yielding a refined particle subset (48k particles). This subset was further refined using Non-uniform Refinement, producing a consensus map.

To improve local resolution, particles were subjected to Local Refinement with a mask applied to the transmembrane domain (TMD), yielding a reconstruction at ~2.9 Å resolution. To further enhance the N-terminal domain (NTD) density, particles were re-centered on the NTD region, followed by re-extraction and Local Refinement with a TMD mask, resulting in an NTD-focused reconstruction at ~4.0 Å resolution.

Model building was performed with initial models generated from homologous AMPAR structures and AlphaFold(*57*) predictions. Models were rigid-body fitted into density maps using UCSF ChimeraX(*58*), followed by iterative manual adjustment in Coot(*59*). Real-space refinement was carried out using PHENIX(*60*), and model geometry was further optimized through iterative cycles of refinement and manual correction. Final models were validated using MolProbity. Figures were prepared using UCSF ChimeraX and PyMOL(*61*).

**Mass spectrometry**

Pig hippocampal P2 fractions were thawed and processed as described above to obtain solubilized AMPAR complexes. Hippocampal P2 pellets were divided into three independent groups, each corresponding to one biological replicate. For each group, 4 mL of P2 pellet was resuspended in 40 mL ice-cold water and incubated at 4 °C for 15 min with gentle rotation. Samples were centrifuged at 23,000 × g for 30 min at 4 °C (Ti-45 rotor), and the resulting pellets were resuspended in lysis buffer containing 150 mM NaCl, 20 mM Tris-HCl (pH 8.0), 1% digitonin, 100 μM PMSF, and protease inhibitor cocktail. Solubilization was carried out for 2 h at 4 °C with gentle rotation, followed by ultracentrifugation at 150,000 × g for 45 min at 4 °C (Ti-45 rotor) to remove insoluble material.

For each replicate, the clarified supernatant was divided into two equal fractions corresponding to antibody-positive and negative control conditions. Each fraction was pre-cleared by incubation with 250 μL protein G bead slurry for 1 h at 4 °C with rotation, followed by centrifugation at 500 × g for 5 min to remove the beads.

For immunoprecipitation, 10 μg of anti-GluA2 C-terminal antibody (mouse monoclonal against epitope CVAKNAQNINPSSSQ produced by GenScript) was added to the antibody-positive samples, while the same volume of lysis buffer was added to the negative controls. Samples were incubated overnight at 4 °C with gentle rotation.

On the following day, 50 μL of protein G bead slurry was added to each sample and incubated for 4 h at 4 °C with rotation. Beads were then pelleted by centrifugation at 500 × g for 1 min, and the flow-through (unbound fraction) was collected. The beads were washed three times with 1 mL wash buffer containing 150 mM NaCl, 20 mM Tris-HCl (pH 8.0), and 0.01% GDN. After removal of excess buffer, the beads were snap-frozen in liquid nitrogen and submitted for mass spectrometry analysis.

#### Quantification of proteins

The raw files were analysed by MaxQuant (version 2.4.2.0) using standard settings, the Label-Free Quantification (LFQ) and intensity-Based Absolute Quantification (iBAQ) options were selected. Spectra were submitted to database search against protein sequences from UP000008227_9823_sus scrofa (downloaded on July 2024) with the following parameters: up to two trypsin missed cleavage sites were allowed; carbamidomethylation of cysteine as fixed modification; and oxidation of methionine and acetylation N-terminal protein set as variable modifications.

Following the approach outlined in, abundances normalised to the AMPAR tetramer were obtained by by multiplying each subunit's iBAQ value by 4 and dividing by the summed iBAQ abundances of GluA1–4.

To obtain protein abundance for statistical testing, peptide-level LFQ intensities were processed in R (v.4.5.2) using QFeatures (v.1.20.0) and biomasslmb (v.0.0.5) R packages. Peptides were filtered to remove matches to contaminants and those with Q-value > 0.01 and intensities were log2-transformed. Peptides quantifyied in fewer than 2 out of 6 samples were removed. Remaining peptides were summarised to protein-level abundances using MsCoreUtils::robustSummary. In total, 209 proteins were quantified. Missing values were imputed exclusively in control samples, and only where all three replicates had missing values. Imputation was performed using MsCoreUtils::impute_minProb, with default settings.

#### Statistical testing

Statistical testing to compare GluA2 IP samples to control samples at each time point was performed with limma (v.3.66.0) using the treat function, setting a minimum relevant fold change of 1.1 as the null hypothesis, using the trend=TRUE argument to account for a mean–variance trend in the prior variance estimation". P values were adjusted using the Benjamini–Hochberg FDR procedure and a threshold of 0.01 (1% FDR) used to identify statistically significant differences.

**Comparison of PRRT1 N-terminal deletion and wild-type expression levels**

HEK293 cells were transfected at a DNA ratio of GluA1:CNIH2:PRRT1-HA = 1:2:2 and harvested 24-26 h post-transfection. All constructs were expressed from PRK5, except CNIH2 from IRES with mCherry. PRRT1 was C-terminally HA tagged for comparison between wild-type and N-terminal deletion.

Cells were collected by centrifugation at 3,000 × g for 10 min at 4 °C and lysed in buffer containing 150 mM NaCl, 20 mM Tris-HCl (pH 8.0), 1% digitonin, protease inhibitors, and phosphatase inhibitor cocktail (PhosSTOP, Roche, Cat# 4906845001) for 2 h at 4 °C. Lysates were clarified by centrifugation at 21,000 × g for 30 min at 4 °C, and the resulting supernatants were analysed by western blotting to compare expression levels of PRRT1 wild-type and N-terminal deletion constructs.

**Immunoprecipitation of phosphorylated and total GluA1-containing AMPAR complexes**

Forebrains from three adult wild-type mice (biological triplicates) were dissected following removal of the cerebellum and olfactory bulbs and homogenized in ice-cold homogenization buffer containing 320 mM sucrose, 10 mM HEPES (pH 7.3), 100 μM PMSF, protease inhibitors, and phosphatase inhibitor cocktails. Homogenates were subjected to sequential centrifugation at 600 × g and 900 × g for 10 min at 4 °C to remove debris and nuclei. The resulting supernatant was centrifuged at 12,000 × g for 30 min at 4 °C to obtain the crude synaptosomal membrane fraction (P2).

P2 membranes were solubilized in lysis buffer containing 1% digitonin, 150 mM NaCl, 20 mM Tris-HCl (pH 8.0), protease inhibitors, and phosphatase inhibitor cocktails for 2 h at 4 °C with gentle rotation. Insoluble material was removed by ultracentrifugation at 150,000 × g for 30 min at 4 °C. The clarified lysate was pre-cleared with Protein A agarose beads for 30 min at 4 °C.

For immunoprecipitation, aliquots of the pre-cleared lysate were incubated with either anti-GluA1 C-terminal antibody (A samples) or phospho-GluA1 Ser831-specific antibody (P samples) for 2 h at 4 °C, followed by capture with Protein A agarose beads for an additional 2 h. Following collection of the phospho-GluA1 immunoprecipitates, the corresponding flow-through fractions were subjected to a second round of immunoprecipitation using the anti-GluA1 C-terminal antibody to isolate the remaining non-phosphorylated GluA1-containing complexes (PA samples).

Immunocomplexes were washed three times with wash buffer supplemented with 0.01% GDN and eluted for SDS–PAGE analysis. Proteins recovered from the A, P, and PA immunoprecipitations were analysed by western blotting using the indicated antibodies. All experiments were performed in biological triplicate.

**Fractionation of hippocampal tissue and Western blot analysis**

Hippocampal tissue was weighed and homogenized in ice-cold homogenization buffer at a 1:10 (w/v) ratio (e.g., 100 mg tissue in 900 μL buffer). Tissue was first homogenized using a blender until fully dissociated, followed by 10 strokes with a Potter–Elvehjem homogenizer. An aliquot of the total homogenate (H) was collected for analysis.

The homogenate was centrifuged at 1,000 × g for 10 min at 4 °C to obtain the crude nuclear fraction (P1) and supernatant (S1). The P1 pellet was washed three times by resuspension in homogenization buffer (1 mL per 200 μL pellet), followed by centrifugation at 1,000 × g for 10 min each time, yielding a washed P1 fraction (P1'). Samples of P1' and S1 were collected.

The S1 fraction was carefully collected and centrifuged at 12,000 × g for 20 min at 4 °C to generate the crude membrane fraction (P2) and cytosolic fraction (S2). Samples of P2 and S2 were collected. The P2 pellet was solubilized in 0.5% Triton X-100 for 15 min on ice and subsequently centrifuged at 150,000 × g for 30 min at 4 °C. This step yielded the postsynaptic density–enriched pellet (PSD/P3) and supernatant (S3), both of which were sampled.

All fractions were mixed 1:1 with 1% RIPA buffer prior to the addition of SDS loading buffer. Samples were resolved on 4–12% Bis-Tris SDS–PAGE gels and transferred onto PVDF membranes. Western blotting was performed using primary antibodies against GluA1, PRRT1, PSD95, Rab5, and Bassoon, followed by HRP-conjugated secondary antibodies.

**DNA constructs and cell culture**

cDNAs encoding rat GluA1 (flip isoform) and rat GluA3R439G (flip isoform) were subcloned into PRK5 vectors. The tandem construct GluA2_TARP-γ8 (flip isoform, edited at Q/R site) was used for heteromeric GluA1/GluA2 recordings. Mutagenesis was performed by IVA cloning as described previously(*62*). cDNAs encoding for CNIH2, TARP-γ8 and PRRT1 (all from rat) were subcloned in IRES vectors. For the N terminal deletion of PRRT1 construct, amino acids from 6 to 206 were deleted. For immunostaining and western blot, an HA epitope was inserted at the end of PRRT1 C terminal with a linker. For organotypic slices, all constructs were expressed from the PRK5 vector. The truncated CaMKII construct (tCaMKII, 1-290 amino acids) fused with EGFP was a gift from José Esteban, and has been described previously(*9*).

HEK293T cells (ATCC CRL-11268, RRID: CVCL_1926; STR-authenticated and mycoplasma-free) were maintained in DMEM supplemented with 10% FBS and penicillin/streptomycin at 37 °C and 5% $CO_2$. Cells were transfected with 1 µg total DNA using Turbofect (Thermofisher) at an AMPAR:CNIH2:PRRT1 ratio of 1:2:2. To limit AMPAR-mediated toxicity, 30 µM NBQX was included during transfection. Recordings and immunostaining were performed 24–48 h post-transfection.

**Electrophysiology (HEK cell recordings)**

Patch pipettes (2-4 MΩ for whole-cell; 6-12 MΩ for outside-out) were fabricated from borosilicate glass and fire-polished. The internal solution contained (mM): 120 CsF, 10 CsCl, 10 EGTA, 10 HEPES, 2 $Na_2$ATP, and 0.1 spermine (pH 7.3, CsOH). The external solution contained (mM): 145 NaCl, 3 KCl, 2 $CaCl_2$, 1 $MgCl_2$, 10 glucose, and 10 HEPES (pH 7.4, NaOH).

Recordings were obtained using an Axopatch 700B amplifier, filtered at 10 kHz, digitized at 100 kHz, and analysed in pClamp 11.2. Cells were plated on poly-L-lysine-coated coverslips on the day of recording. Rapid solution exchange was achieved using a piezo-driven theta-glass application system (20-80% rise time: ~120 μs for outside-out). Membrane patches were held at -60 mV (uncorrected for an 8.5 mV junction potential). Desensitization (200 ms application of 10 mM glutamate) and deactivation (1 ms pulse of 10 mM glutamate) time constants were determined by fitting the current decay from 90% of peak to baseline or steady state with single- or double-exponential functions using the Chebyshev algorithm (Clampfit 11.2, Molecular Devices). For double-exponential fits, the weighted time constant ($\tau_{w,des}$) was calculated as $\tau_{w,des} = \tau_f(A_f/(A_f + A_s)) + \tau_s(A_s/(A_f + A_s))$, where $\tau_f$ and $\tau_s$ are the fast and slow time constants, and $A_f$ and $A_s$ are their respective amplitudes.

**Animals**

All mouse brain tissue used was extracted from C57BL/6J Ola mice (RRID: MGI:3691859) (age 4-6 weeks for all experimental use except for organotypic brain slice). Mouse brain tissue for expansion microscopy was prepared using transcardial fixative perfusion method. Adult C57BL/6J Ola mice were deeply anesthetized with sodium pentobarbital. They were then transcardially perfused first with PBS followed by 4% paraformaldehyde (PFA) in PBS. Brains were collected and post-fixed for 20-24 h at 4 °C in the same fixative solution, then washed in PBS before slicing.

All procedures were carried out under PPL PP5747704 in accordance with UK Home Office regulations and licensed under the UK Animals (Scientific Procedures) Act of 1986, following local ethical approval. All animals were housed with unlimited access to food and water on a 12 h–12 h light–dark cycle at room temperature (20–22 °C) and 45–65% humidity. Sample size was based on previous experience with this sort of experiment and there was no randomization or blinding.

**Brain slice preparation**

For organotypic slice culture(*63*), hippocampi from (postnatal) P6–7 mice were dissected in ice-cold Gey's balanced salt solution containing (mM): 175 sucrose, 150 NaCl, 2.5 KCl,

0.85 $NaH_2PO_4$, 0.66 $KH_2PO_4$, 2.7 $NaHCO_3$, 0.28 $MgSO_4$, 2 $MgCl_2$, 0.5 $CaCl_2$ and 25 d-glucose at pH 7.3. Hippocampi were cut using a McIlwain tissue chopper into 300 μm slices that were then grown on Millicell cell culture inserts (Merck) in culture medium (78.5% MEM, 15% heat-inactivated horse serum, 2% B27+ supplement, 2.5% 1 M HEPES, 1.5% 0.2 M L-Glutamine, 0.5% 0.05 M ascorbic acid, 1 mM $CaCl_2$, 1 mM $MgSO_4$) at 37 °C and 5% CO2. Cultures were transfected at 7-9 DIV by single-cell electroporation. Recordings were performed 3-4 days after transfection.

For expansion microscopy, perfusion-fixed mouse brains were sliced coronally at a 50-μm thickness using a vibratome (Leica VT 1200 S) and stored at 4 °C in PBS before expansion.

**Single-cell electroporation**

Single cells from the CA1 region of organotypic hippocampal slices were transfected using an adapted version of the method described previously(*64, 65*). The transfection ratio of AMPAR:PRRT1:tCaMKII DNA plasmids was 2:2:3, of AMPAR or PRRT1:tCaMKII was 2:3, of AMPAR:PRRT1 was 1:1, and PN1-eGFP was added to aid visualisation. DNA plasmid mixture was diluted to 33 ng μl−1 in intracellular solution containing (mM): 125 KGlu, 20 KCl, 4 $MgCl_2$, 10 HEPES, 4 Na2-ATP, 0.3 Na-GTP, 0.2 EGTA and back-filled into borosilicate microelectrode pipettes (4–8 MΩ). Slices were placed into the recording chamber sterilized with 70% ethanol and filled with HEPES-based artificial cerebrospinal fluid (aCSF) containing (mM): 140 NaCl, 3.5 KCl, 1 $MgCl_2$, 2.5 $CaCl_2$, 10 HEPES, 10 glucose, 1 Na-pyruvate, 2 $NaHCO_3$. Cells were briefly kept in cell-attached mode and DNA was introduced with a short burst of current pulses (60 pulses at 200 Hz). Slices were returned to incubation in their original culture medium supplemented with 5 μg $ml^{-1}$ gentamycin until recording.

**Slice electrophysiology**

Transfected hippocampal slice cultures were used for electrophysiological recording 3–4 days after single-cell electroporation. Slices were perfused with aCSF containing (mM): 10 d-glucose, 26.4 $NaH_2CO_3$, 126 NaCl, 1.25 $NaH_2PO_4$, 3 KCl, 4 $MgSO_4$, 4 $CaCl_2$ saturated with

95% $O_2$/5% $CO_2$. Then, 100 μM D-APV, 1 μM SR-95531 and 2 μM 2-chloroadenosine were added to the aCSF for synaptic recording. Borosilicate pipettes (4–6 MΩ) were filled with intracellular solution containing (mM): 135 $CsMeSO_4$, 4 NaCl, 2 $MgCl_2$, 10 HEPES, 4 $Na_2$-ATP, 0.4 Na-GTP, 0.15 spermine, 0.6 EGTA, 0.1 $CaCl_2$, adjusted to pH 7.3 with CsOH. All whole-cell recordings were dual, involving simultaneous recording of a neighbouring pair of transfected (GFP positive) and untransfected cells. EPSCs were evoked by Schaffer collateral stimulation at 0.2 Hz in the stratum radiatum at the CA3–CA1 border. EPSCs represent an average of at least 20 sweeps. Recordings were excluded if the series resistance exceeded 20 MΩ or varied by more than 20%. Recordings were made using pClamp10 (Molecular Devices) with a Multiclamp 700B amplifier (Axon Instruments), digitized using a Digidata 1440A (Axon Instruments), and analysed in pClamp 11.2.

**Tissue expansion with Magnify**

Magnify gelling solution was prepared with 34% (w/v) sodium acrylate (lab preparation), 10% (w/v) acrylamide, 4% (w/v) N,N-dimethylacrylamide acid (DMAA), 1% (w/v) NaCl and 0.01% (w/v) bis-acrylamide in PBS (all chemicals from Sigma Aldrich and Thermo Fisher Scientific) and kept at 4°C. Immediately before incubation with the sample, the following chemicals were added: ammonium persulfate (APS; 0.2%, w/v), N,N,N',N'-tetramethyl-ethane-1,2-diamine (TEMED; 0.2%, w/v), 2,2,6,6-tetramethylpiperidin-1-oxyl (TEMPO; 0.001%, w/v) and methacrolein (0.2%, v/v); all chemicals from Sigma Aldrich and Thermo Fisher Scientific. Brain slices were incubated with the gelling solution for 30 min at 4°C and then transferred to gelling chambers for 2 h incubation in a humidified container at 37 °C to complete gelation.

After gelation, brain slices embedded in gel were removed from the chamber and trimmed to only preserve the hippocampus region. Samples were then incubated in homogenization buffer (200 mM SDS, 8 M Urea, 25 mM EDTA, 2× PBS, pH 7.5 at RT; all chemicals from Sigma Aldrich) for 15–20 h at 80°C. Homogenized samples were then washed two times with 1% decaethylene glycol mono-dodecyl ether (DGME; Sigma Aldrich) in PBS at 60°C, followed by at least three 20 min washes in PBS at room temperature.

Samples were then placed into an excess volume of ddH2O (at least 10-times the final gel volume) for expansion. The ddH2O was replaced 4-5 times, with 10-15 min per expansion step, to reach the final expansion factor of ~7–8×.

Expanded hippocampal samples were stabilized by re-embedding with 6% (w/v) acrylamide, 0.15% (w/v) bis-acrylamide, and 5 mM Tris-HCl (pH 8.0) in ddH2O and kept on ice. Immediately before incubation with the sample, the following chemicals were added: APS (0.1%, w/v) and TEMED (0.1%, w/v). Samples were incubated with the re-embedding solution for 30 min at 4°C and then transferred to gelling chambers for 2 h incubation in a humidified container at 37 °C to complete stabilization. Hippocampal samples were then removed from the chamber, trimmed as necessary, and stored at 4°C in PBS before immunostaining.

**Immunostaining**

For HEK cells, cells were plated onto poly-l-lysine-treated glass coverslips after 24 h of transfection and used for immunostaining after 48 h of transfection. Cells were fixed in phosphate-buffered saline (PBS) containing 4% paraformaldehyde and 4% sucrose at 4 °C for 20 min, washed in PBS and treated with blocking solution containing 5% bovine serum albumin (Fisher Bioreagents, Cat# BP1605-100) and 10% normal goat serum (Sigma-Aldrich, Cat# G9023) in PBS. For surface staining, cells were incubated with anti-HA tag primary antibody solutions prepared in PBS supplemented with 5% BSA and 10% normal goat serum for 2-4 h at room temperature, and washed in PBS after incubation. For permeabilised staining, cells were then treated in 0.1% Triton-X 100 added blocking solution and sequentially incubated with anti-GluA1 primary antibody for 2-4 h and secondary antibodies for 1-2 h. Coverslips were mounted in ProLong Glass antifade Mountant (Invitrogen, Cat# P36980) and left to cure for 24 h at room temperature before confocal imaging.

For post-expansion immunostaining, mouse hippocampal samples were blocked in blocking solution containing PBS, 0.2% Triton-X 100, 5% BSA, and 10% donkey serum for 1 h. Samples were then incubated with primary antibodies at 4°C for 40–48 h and secondary antibodies at 4°C for 15–20 h, with three 30 min PBS washes after each incubation. Immunostaining was followed by 1 h DAPI (1:1000 from 1 mg/ml stock) staining followed by three 10 min PBS washes before confocal imaging.

**Antibodies**

For immunostaining, the following primary antibodies were used: mouse anti-HA.11 epitope tag (Biolegend, Cat# 901513, RRID:AB_2565335, clone 16B12, 1:500), anti-glutamate receptor 1 (AMPA subtype) antibody (Abcam, Cat# ab31232, RRID: AB_2113447, 1:500), GluA1 (AMPA1) antibody (Synaptic System, Cat# 182 011, 1:500), SynDIG4/PRRT1 antibody (Proteintech, Cat# 17261-1-AP, RRID: AB_2878371, 1:150), Shank2 antibody (Synaptic System, Cat# 162 204, 1:500), Anti-Bassoon antibody (Abcam, Cat# ab82958, RRID: AB_1860018, clone SAP7F407, 1:500). The following secondary antibodies were used: goat anti rabbit IgG conjugated with Alexa Fluor 405 (Invitrogen, Cat# A31556, RRID: AB_221605, 1:300), goat anti guinea pig IgG conjugated with Alexa Fluor 488 (Invitrogen, Cat# A11073, RRID: AB_2534117, 1:300), donkey anti rabbit IgG conjugated with Alexa Fluor 568 (Invitrogen, Cat# A10042, RRID: AB_2534017, 1:300), goat anti mouse IgG conjugated with Alexa Fluor 647 (Invitrogen, Cat# A21235, RRID: AB_2535804, 1:300), donkey anti mouse IgG conjugated with Alexa Fluor 647 (Jackson ImmunoResearch, Cat# 715 605 150, RRID: AB_2340862, 1:300).

For western-blot, the following primary antibodies were used: anti-HA antibody (Abcam, Cat# ab137838, 1:200), anti-glutamate receptor 1 (AMPA subtype) antibody (Abcam, Cat# ab31232, 1:1000), anti-phospho-GluA1-S831 (Sigma-Aldrich, Cat# 04-823, clone N453, 1:1000), SynDIG4/PRRT1 antibody (Proteintech, Cat# 17261-1-AP, 1:500), anti-PSD95 antibody (SYSY, Cat#124002, 1:1000), anti-Rab5 antibody (Abcam, Cat# ab18211, 1:1000), anti-Bassoon antibody (Abcam, Cat# ab82958, clone SAP7F407, 1:500). The following secondary antibodies were used: mouse Anti-Rabbit light chain, HRP conjugate antibody (Sigma-Aldrich, Cat# MAB201P, 1:4000), native Staphylococcus aureus Protein A HRP (Abcam, Cat#ab7456).

**Confocal imaging**

Images were acquired with a Leica TCS SP8 confocal inverted microscope controlled by Leica Application Suite X (LAS X) software and equipped with a 405-nm diode laser, a tuneable pulsed white light laser, a 633-nm HeNe laser and hybrid detectors. Images were acquired using a 20x/0.75NA objective for overview image of organotypic slice, a 63x/1.4NA oil-immersion objective for HEK cells, and a 40x/1.1NA water-immersion objective for expansion microscopy. Alexa Fluor 405, Alexa Fluor 488, Alexa Fluor 568 and Alexa Fluor 647 were excited at 405 nm, 496 nm, 587 nm and 633 nm, respectively. Z-stacks at 0.4-μm spacing were acquired with laser intensities adjusted for each biological replicates (independent expansion and staining trials, each from different slices from different adult mice). Images were processed and pseudo-coloured in Fiji (ImageJ) for display purposes. 3D multiple spot colocalization was analysed using a custom-written macro in Fiji. In brief, spots in 3D stacks were detected by pre-defined threshold in each channel and sub-pixel localization was achieved by a block-coordinate Gaussian least square curve fitting. Spot colocalization was defined by a distance threshold of 1 μm (expanded scale) and counted without a pre-defined reference mark. The final counts of all different types of colocalization events across all channels were aggregated, corrected for lower-cardinality sets, and normalized to set cardinality. Parameter settings were manually checked and adjusted with a testing image and applied throughout the analysis. 5,000–8,000 total spots were found in each image stack.

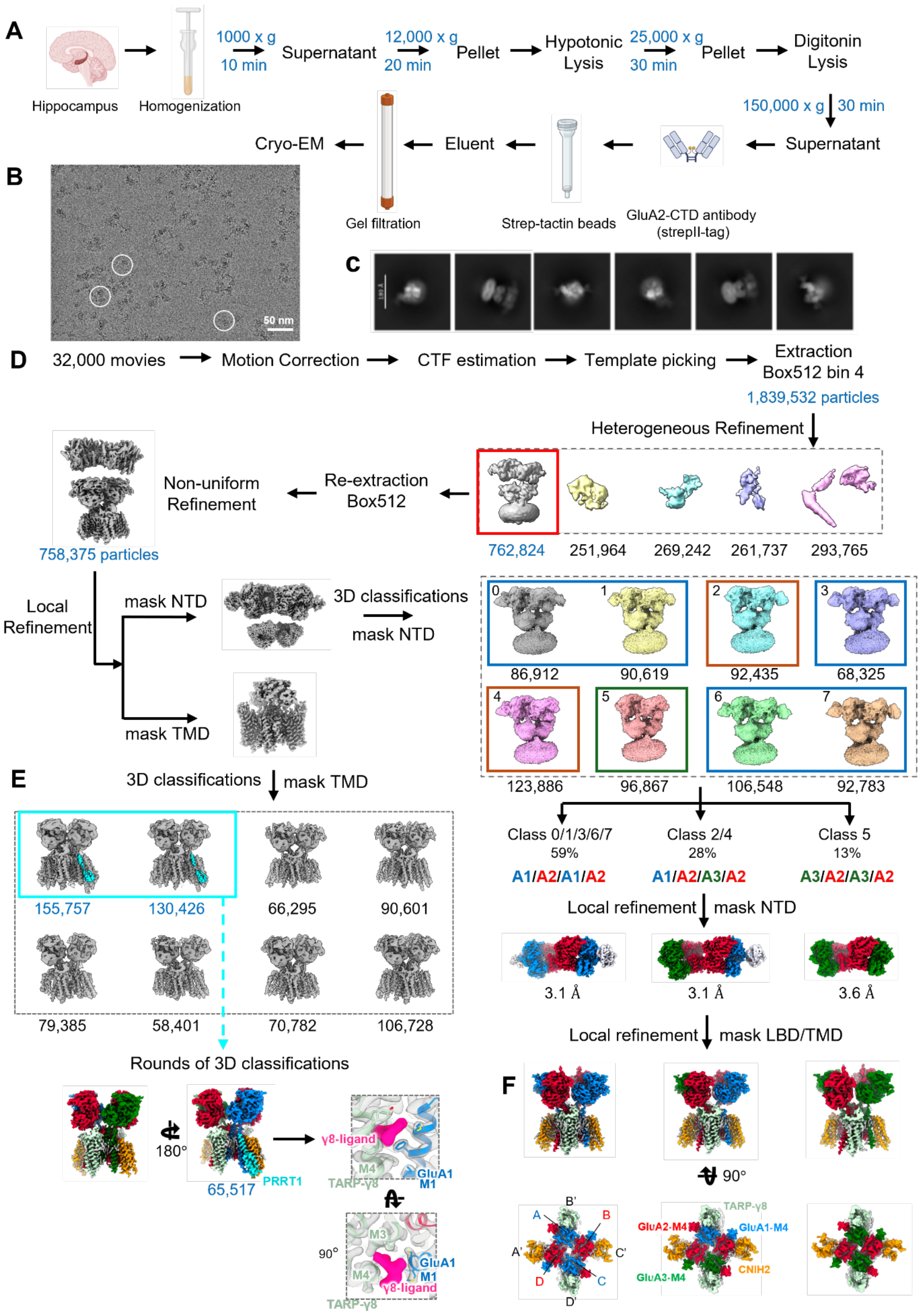
A
Hippocampus
Homogenization
1000 x g 10 min
Supernatant
12,000 x g 20 min
Pellet
Hypotonic Lysis
25,000 x g 30 min
Pellet
Digitonin Lysis
150,000 x g 30 min
Supernatant
GluA2-CTD antibody (strepII-tag)
Strep-tactin beads
Eluent
Gel filtration
Cryo-EM
B
50 nm
C
D
32,000 movies
Motion Correction
CTF estimation
Template picking
Extraction Box512 bin 4
1,839,532 particles
Heterogeneous Refinement
762,824
251,964
269,242
261,737
293,765
Re-extraction Box512
Non-uniform Refinement
758,375 particles
Local Refinement
mask NTD
mask TMD
3D classifications
mask NTD
86,912
90,619
92,435
68,325
123,886
96,867
106,548
92,783
Class 0/1/3/6/7
59%
A1/A2/A1/A2
Class 2/4
28%
A1/A2/A3/A2
Class 5
13%
A3/A2/A3/A2
Local refinement
mask NTD
3.1 Å
3.1 Å
3.6 Å
Local refinement
mask LBD/TMD
E
3D classifications
mask TMD
155,757
130,426
66,295
90,601
79,385
58,401
70,782
106,728
Rounds of 3D classifications
180°
65,517
PRRT1
γ8-ligand
M4
TARP-γ8
GluA1
M1
90°
M3
M4
GluA1
M1
γ8-ligand
TARP-γ8
F
90°
A
B
B'
A'
C'
D
C
D'
GluA2-M4
GluA1-M4
TARP-γ8
CNIH2
GluA3-M4

480

**Fig. S1. Purification and cryo-EM analysis workflow of native GluA2 AMPAR complexes**

**A**, Workflow for purification of native AMPAR complexes from pig hippocampus using a GluA2-CTD antibody. Hippocampal tissue was homogenized and subjected to differential centrifugation to isolate the P2 synaptosome fraction. Following hypotonic lysis and digitonin solubilization, clarified supernatants were incubated with Strep-Tactin beads loaded with Strep-tagged GluA2-CTD antibody. Eluted complexes were further purified by gel filtration prior to cryo-EM analysis.

**B**, Representative cryo-EM micrograph of purified native AMPAR particles embedded in vitreous ice. Example particles are highlighted with white circles.

**C**, Representative 2D class averages showing characteristic views of native AMPAR complexes.

**D**, Cryo-EM image processing workflow for native AMPAR datasets. A total of 32,000 movies were collected and processed through motion correction, CTF estimation, template-based particle picking, heterogeneous refinement, and iterative 3D classification. Three principal receptor assemblies were identified: A1/A2/A1/A2 (59%), A1/A2/A3/A2 (28%), and A3/A2/A3/A2 (13%). Final local refinements yielded reconstructions at 3.1 Å, 3.1 Å, and 3.6 Å resolution, respectively.

**E**, Focused classification of the transmembrane domain identified a subset of particles containing additional PRRT1 density (cyan). Iterative rounds of masked 3D classification enriched PRRT1-containing particles, yielding a final subset of 65,517 particles with well-resolved PRRT1-associated density. The enlarged view of the γ8–ligand interface is indicated by the black dashed box.

**F**, Further local refinement of the three principal receptor assemblies using a masked LBD/TMD region showed that all classes are assembled with two CNIH2 and two TARP-γ8 subunits. Bottom views of the corresponding LBD/TMD classes are shown. The rotation angle and viewing direction are indicated.

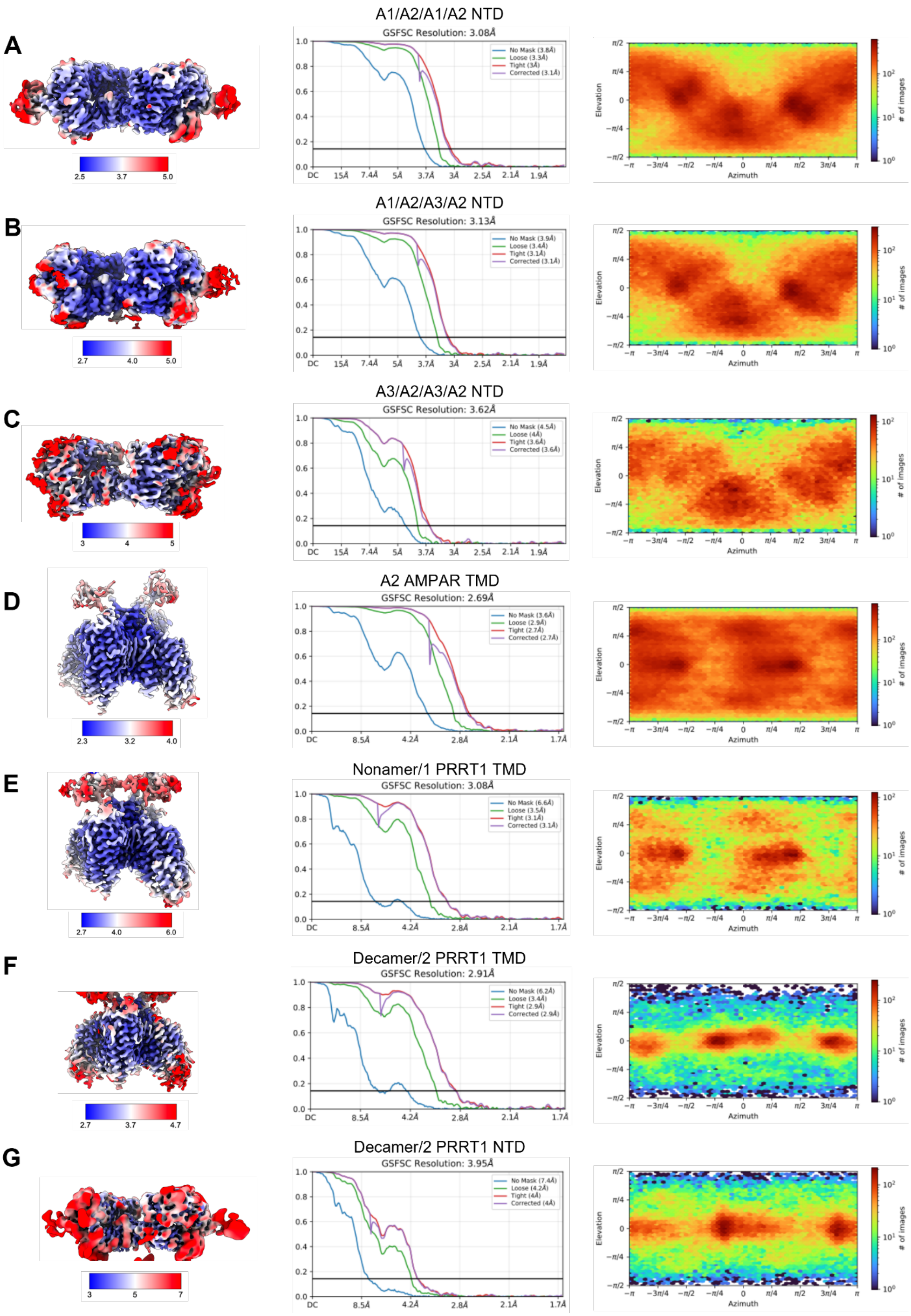
A
A1/A2/A1/A2 NTD
GSFSC Resolution: 3.08Å
No Mask (3.8Å)
Loose (3.3Å)
Tight (3Å)
Corrected (3.1Å)
B
A1/A2/A3/A2 NTD
GSFSC Resolution: 3.13Å
No Mask (3.9Å)
Loose (3.4Å)
Tight (3.1Å)
Corrected (3.1Å)
C
A3/A2/A3/A2 NTD
GSFSC Resolution: 3.62Å
No Mask (4.5Å)
Loose (4Å)
Tight (3.6Å)
Corrected (3.6Å)
D
A2 AMPAR TMD
GSFSC Resolution: 2.69Å
No Mask (3.6Å)
Loose (2.9Å)
Tight (2.7Å)
Corrected (2.7Å)
E
Nonamer/1 PRRT1 TMD
GSFSC Resolution: 3.08Å
No Mask (6.6Å)
Loose (3.5Å)
Tight (3.1Å)
Corrected (3.1Å)
F
Decamer/2 PRRT1 TMD
GSFSC Resolution: 2.91Å
No Mask (6.2Å)
Loose (3.4Å)
Tight (2.9Å)
Corrected (2.9Å)
G
Decamer/2 PRRT1 NTD
GSFSC Resolution: 3.95Å
No Mask (7.4Å)
Loose (4.2Å)
Tight (4Å)
Corrected (4Å)
Elevation
Azimuth
### of images

**Fig. S2. Structural and biochemical characterization of PRRT1-containing native AMPAR complexes**

**A-G**, Left: local resolution and overall map quality of the GluA2 and PRRT1 containing AMPAR receptors (A-G). Middle: local resolution maps were computed for each voxel, with resolutions determined at Fourier shell correlation (FSC) = 0.143. Gold standard Fourier shell correlation (GSFSC) curves for each map are shown in the middle lane of each panel. Black line is FSC = 0.143. The Y axis is FSC, X is resolution in Å.

Right: Heat maps of particle orientation distribution for each structure are shown on the right lane of each panel.

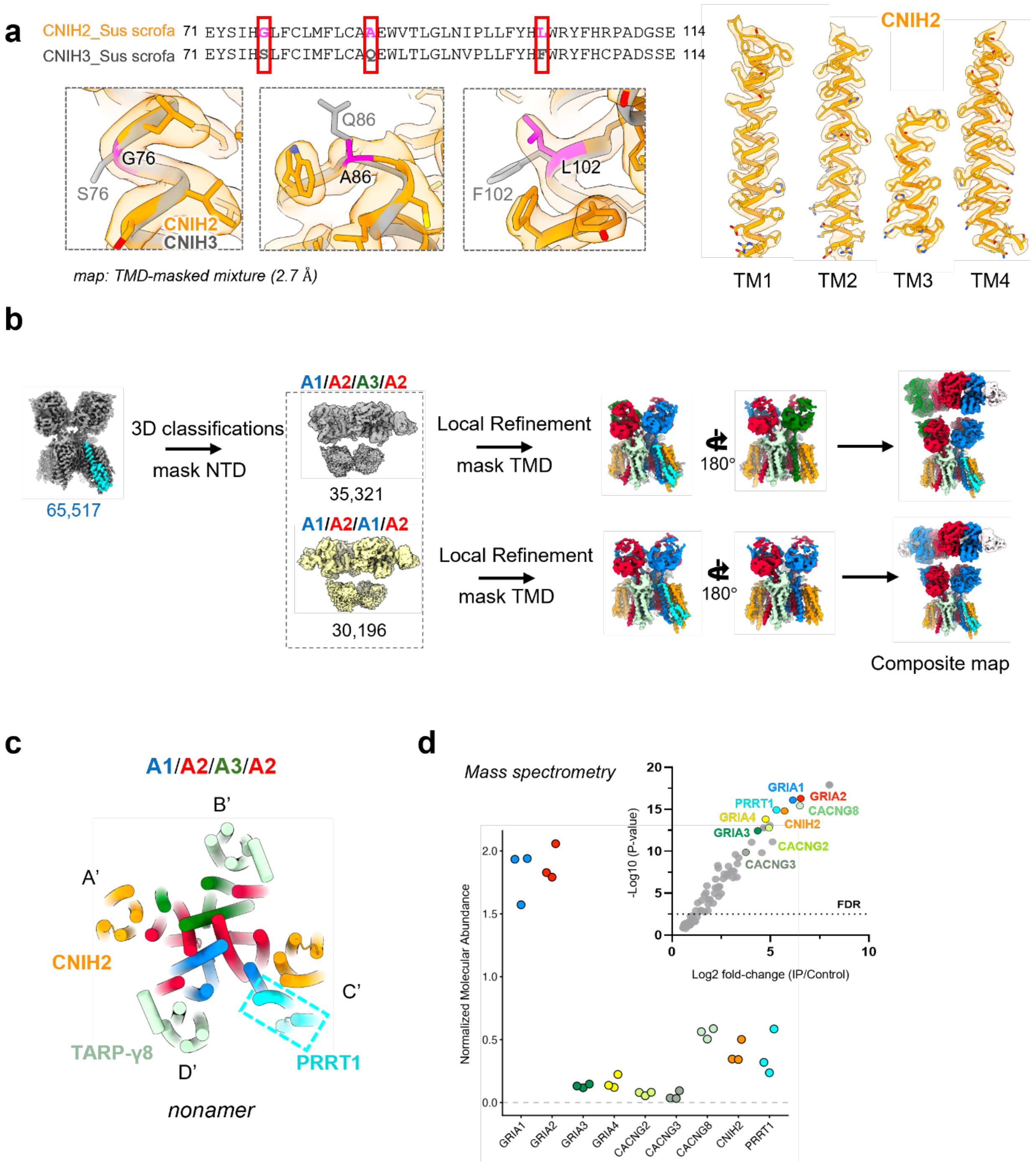


**Fig. S3. Structural and biochemical characterization of native AMPAR complexes**

**A**, Left, sequence alignment of Sus scrofa CNIH2 and CNIH3 spanning residues 71–114. Residues that differ between the two proteins are highlighted in red boxes. Cryo-EM densities corresponding to the indicated positions support assignment of the auxiliary subunit as CNIH2 in the TMD-masked reconstruction (2.7 Å). Right, model-to-map fit of the TM1–TM4 helices of CNIH2.

**B**, Cryo-EM processing workflow used to separate PRRT1-containing receptor assemblies into distinct subunit compositions. Focused 3D classification using an NTD mask separated particles into A1/A2/A3/A2 and A1/A2/A1/A2 populations, followed by local refinement using a TMD mask. Composite maps were generated from the locally refined reconstructions.

**C**, Top views of the TMD models of the nonameric complexes showing subunit arrangement (A'–D') and auxiliary-subunit occupancy. PRRT1 (cyan, dashed boxes) occupies a single auxiliary-subunit binding site alongside CNIH2.

**D**, Quantitative mass spectrometry analysis of native GluA2 pulldown samples. Top right, volcano plot showing protein enrichment ($\log_2$ fold change) versus statistical significance ($-\log_{10}$ P value). Selected AMPAR complex components and associated proteins are highlighted and labelled. The dashed horizontal line indicates the 1% false discovery rate (FDR) threshold. Bottom left, stoichiometry analysis of the core AMPAR complex showing the relative abundance of receptor subunits and associated auxiliary proteins.

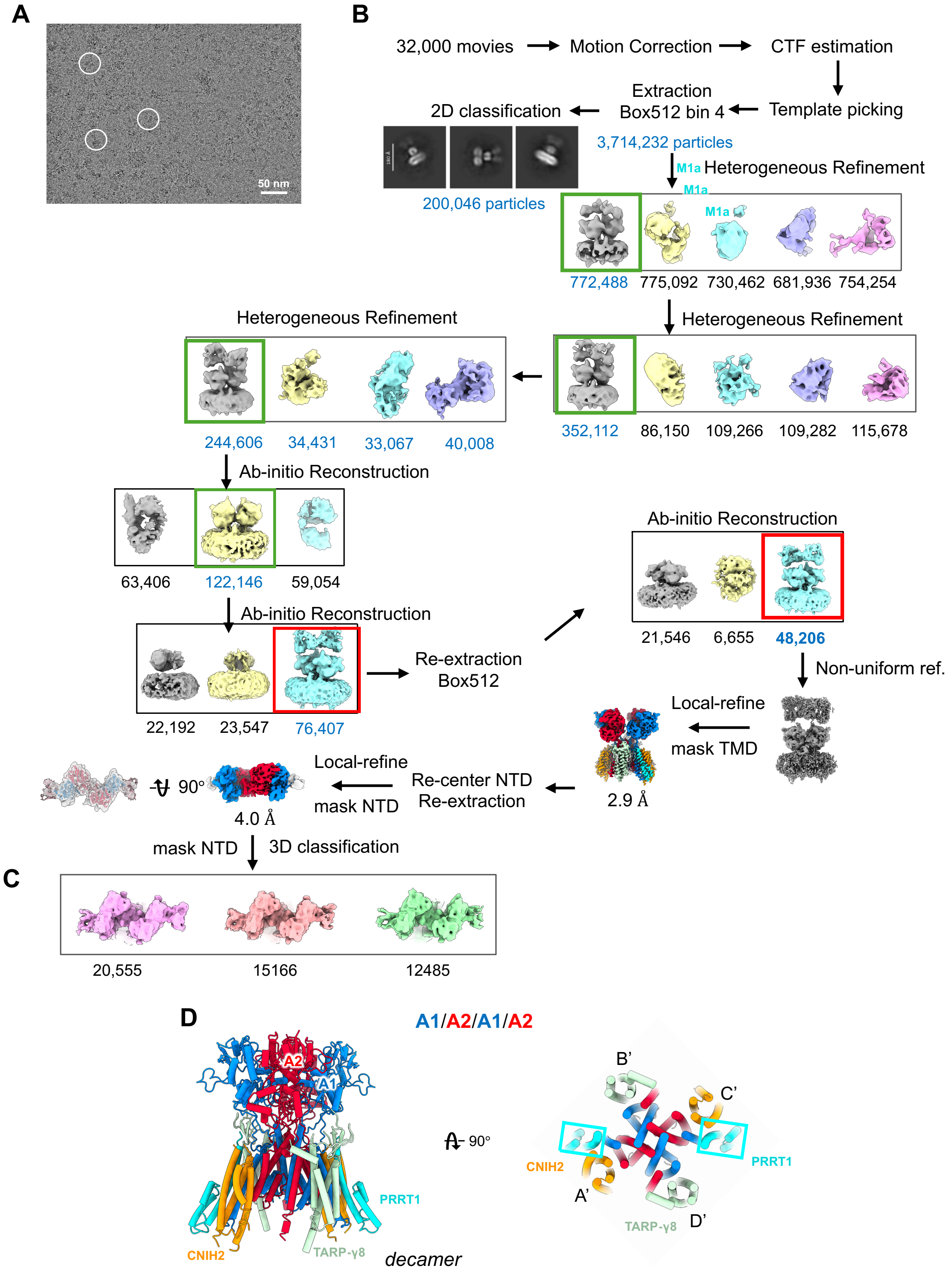
A
50 nm
B
32,000 movies
Motion Correction
CTF estimation
Template picking
Extraction
Box512 bin 4
2D classification
3,714,232 particles
200,046 particles
M1a
Heterogeneous Refinement
772,488
775,092
730,462
681,936
754,254
Heterogeneous Refinement
352,112
86,150
109,266
109,282
115,678
Heterogeneous Refinement
244,606
34,431
33,067
40,008
Ab-initio Reconstruction
63,406
122,146
59,054
Ab-initio Reconstruction
22,192
23,547
76,407
Re-extraction
Box512
Ab-initio Reconstruction
21,546
6,655
48,206
Non-uniform ref.
Local-refine
mask TMD
2.9 Å
Re-center NTD
Re-extraction
Local-refine
mask NTD
90°
4.0 Å
mask NTD
3D classification
C
20,555
15166
12485
D
A1/A2/A1/A2
A2
A1
PRRT1
CNIH2
TARP-γ8
decamer
90°
B'
C'
CNIH2
PRRT1
A'
TARP-γ8
D'

**Fig. S4. Cryo-EM analysis and structural features of PRRT1-containing native AMPAR complexes purified using a PRRT1 N-term antibody**

**A**, Representative cryo-EM micrograph of native hippocampal AMPAR complexes purified using the PRRT1N-term antibody. Example particles are highlighted with white circles.

**B**, Cryo-EM data-processing workflow for PRRT1 N-term pulldown samples. Following motion correction, CTF estimation, template picking, and particle extraction, multiple rounds of heterogeneous refinement and ab-initio reconstruction were performed to separate receptor populations. A PRRT1-containing AMPAR class was refined to 2.9 Å resolution using local refinement with a TMD mask. Subsequent re-centering and local refinement with an NTD mask yielded a 4.0 Å NTD reconstruction. Particle numbers for each processing step are indicated.

**C,** Focused 3D classification of the NTD region identified subclasses showing comparable densities for the GluA1 probe 11B8 at the receptor N-terminal domains.

**D,** Side (left) and top (right) views of the TMD models of the decameric complexes showing subunit arrangement (A'–D') and auxiliary-subunit occupancy. PRRT1 (cyan, dashed boxes) occupies double auxiliary-subunit binding sites alongside CNIH2.

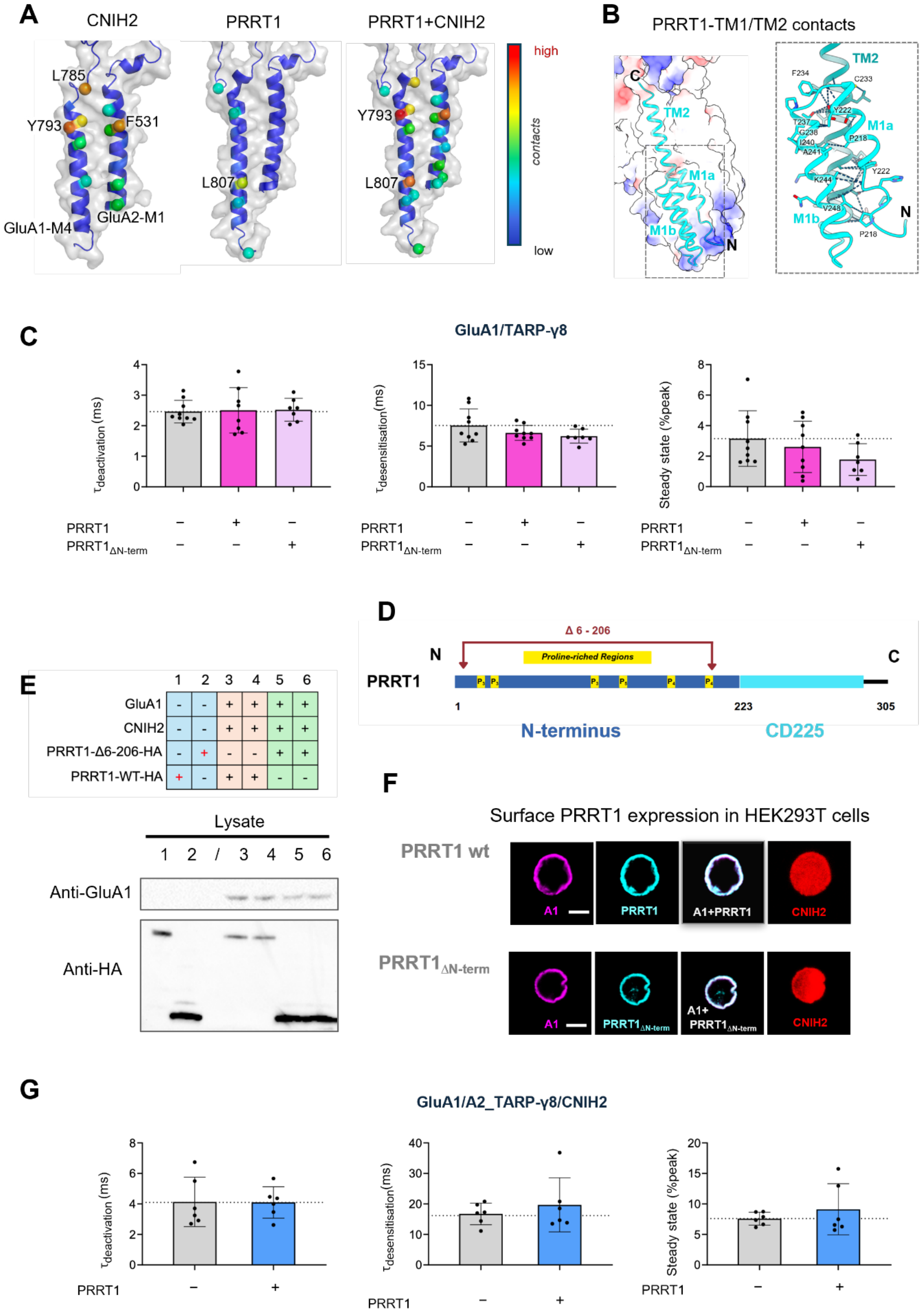
A
CNIH2
PRRT1
PRRT1+CNIH2
L785
Y793
F531
GluA1-M4
GluA2-M1
L807
high
contacts
low
B
PRRT1-TM1/TM2 contacts
TM2
M1a
M1b
C
N
F234
C233
Y222
T237
G238
P218
I240
A241
K244
V248
C
GluA1/TARP-γ8
τdeactivation (ms)
τdesensitisation(ms)
Steady state (%peak)
PRRT1
PRRT1ΔN-term
D
Δ 6 - 206
Proline-rich Regions
N
C
PRRT1
1
223
305
N-terminus
CD225
E
GluA1
CNIH2
PRRT1-Δ6-206-HA
PRRT1-WT-HA
Lysate
Anti-GluA1
Anti-HA
F
Surface PRRT1 expression in HEK293T cells
PRRT1 wt
PRRT1ΔN-term
A1
PRRT1
A1+PRRT1
CNIH2
PRRT1ΔN-term
A1+ PRRT1ΔN-term
G
GluA1/A2_TARP-γ8/CNIH2
τdeactivation (ms)
τdesensitisation(ms)
Steady state (%peak)
PRRT1

**Fig. S5. Structural and functional characterization of PRRT1 interactions with AMPAR complexes**

**A**, Contact maps showing interactions of CNIH2 and PRRT1 within their auxiliary-subunit binding sites, formed by the GluA1 M4 and GluA2 M1 helices. Contacting residues are coloured according to the number of atomic contacts contributing to the interaction (red, high; blue, low). Contacts were calculated using the *ndNeighbors* function in ProDy with a 4.5 Å heavy-atom distance cutoff

**B**, Electrostatic surface map (left, transparent) and structural model of the PRRT1 transmembrane helices TM1 and TM2. Enlarged views (right) highlight side-chain interactions stabilizing the packing between TM1a, TM1b, and TM2. Key interacting residues located within 4 Å are indicated.

**C**, Kinetic parameters of currents evoked by 10 mM glutamate in outside-out patches pulled from HEK293T cells expressing GluA1/TARP-γ8 with or without $PRRT1_{wt}$ constructs. Bar plots show deactivation kinetics during 1 ms glutamate application (left), desensitization kinetics (middle), and steady-state current measured during 200 ms glutamate applications (right) for GluA1/TARP-γ8, GluA1/TARP-γ8 + PRRT1, and GluA1/TARP-γ8 + $PRRT1_{\Delta N\text{-term}}$. n = 8 patches for all conditions. Height of the bar represents mean and whiskers represent standard deviation. The dotted line shows mean kinetic values measured for GluA1/TARP-γ8.

**D**, Schematics of PRRT1 and generation of N-terminal deletion mutants. The PRRT1 N-terminus deletion of 6-206 amino acids (dark red) preserves intact C-terminus (black line), CD225 domain (cyan), and necessary N-terminus (dark blue) for protein synthesis and folding. PRRT1 N-terminus features 5 proline-rich regions (yellow box; P for proline with the number of proline cluster indicated in subscript).

**E,** Western blot analysis of $PRRT1_{\Delta N\text{-term}}$ and $PRRT1_{wt}$ expression. HEK293 cell lysates were collected and analysed by western blotting using anti-GluA1 antibody to detect GluA1 and anti-HA antibody to detect PRRT1. The plasmid combinations used for transfection are indicated in the table above. Groups 3 and 4, and groups 5 and 6, represent biological replicates.

**F**, Representative confocal images of HEK293T cells co-transfected with GluA1 (magenta), PRRT1-HA (cyan), and CNIH2 (red). Surface expression of PRRT1 was confirmed by targeting extracellular C-terminally HA tagged PRRT1 with anti-HA tag antibody before cell permeabilization for GluA1 staining. mCherry was expressed from the same plasmid with CNIH2. Scale bar: 5 μm.

**G**, Kinetic parameters of currents evoked by 10 mM glutamate in outside-out patches pulled from HEK293T cells expressing A1/A2(R)_TARP-γ8/CNIH2 with or without PRRT1 constructs. Bar plots show deactivation kinetics during 1 ms glutamate application (left), desensitization kinetics (middle), and steady-state current measured during 200 ms glutamate applications (right) for A1/A2(R)_TARP-γ8/CNIH2, A1/A2(R)_TARP-γ8/CNIH+ PRRT1, and A1/A2(R)_TARP-γ8/CNIH2 + PRRT1$_{\Delta N\text{-term}}$. n = 6 patches for all conditions. Bars and whiskers as in C.

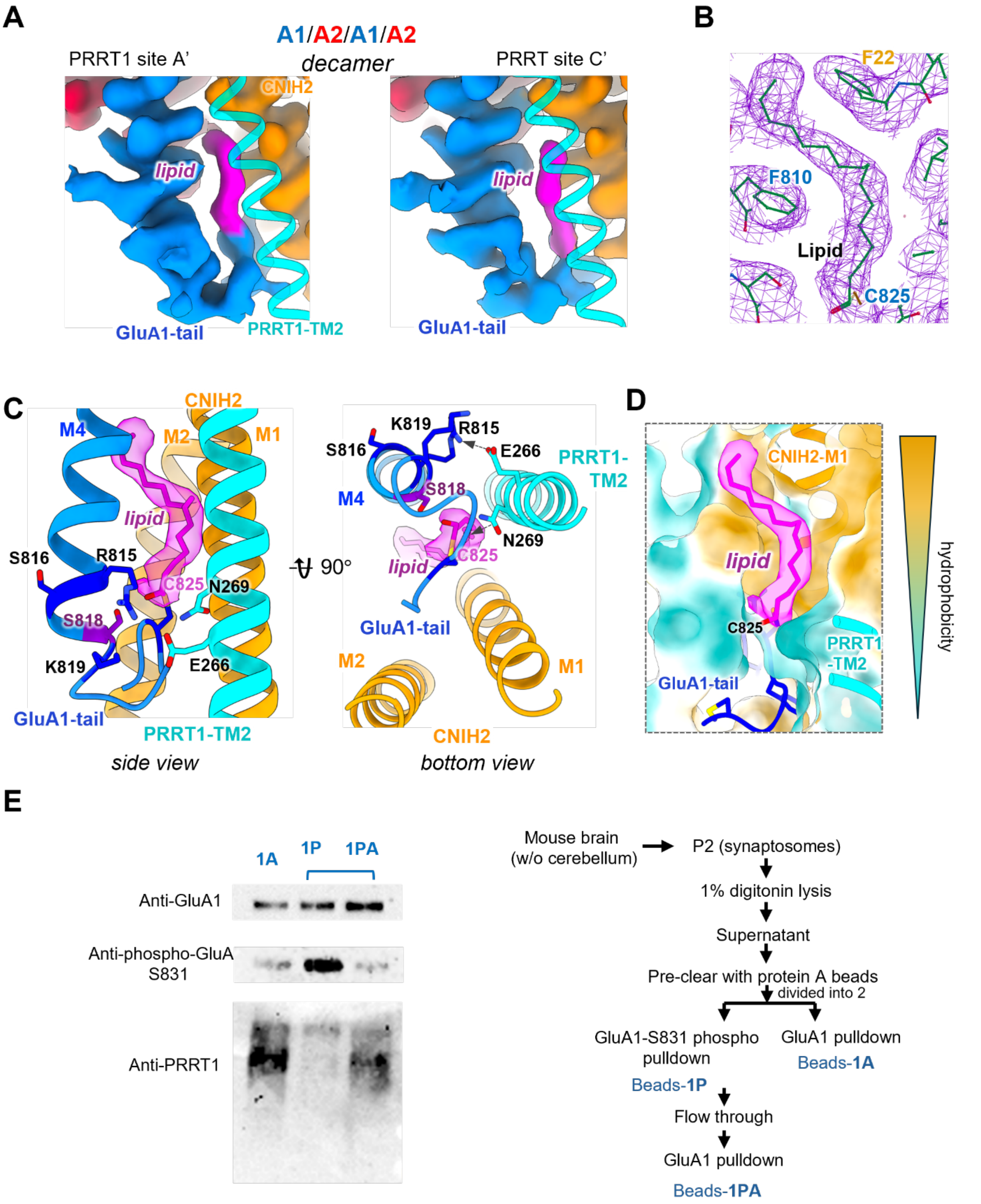


**Fig. S6. Structural characterisation of GluA1-Ctail interaction with M1b helix of PRRT1via C825 lipid**

**A**, Cryo-EM density of the decameric AMPAR complex reconstructed with C1 symmetry, highlighting the dual PRRT1-binding sites. PRRT1 density is shown as a transparent model occupying both sites. On both sides, strong lipid-like density (pink) is observed and is linked to the GluA1 C-terminus.

**B**, Model-to-map fit of lipid-like density associated with GluA1 C825. Surrounding hydrophobic residues are indicated, including F810 of GluA1 and F22 of CNIH2.

**C**, Zoomed-in view of the decameric model illustrating the interaction between the GluA1 M4 region and PRRT1 TM2 in the presence of lipid-like density. Residues R815, S816, S818, and K819 of GluA1 M4 are shown. PRRT1 residues E266 and N269 are indicated. Hydrogen bonding interactions are observed between GluA1 R815/K819 and PRRT1 E266/N269. We note that Ser818, a PKC substrate, is inaccessible to phosphorylation resulting from PRRT1 engagement of the GluA1 C-tail.

**D**, The lipid-like density is embedded within a hydrophobic environment formed by GluA1 M4, PRRT1 TM2, and CNIH2 TM1. The model includes the GluA1 C-terminal together with partial PRRT1 TM2 and CNIH2 TM1.

**E,** Immunoprecipitation of phosphorylated and total GluA1-containing AMPAR complexes. Left, western blot analysis of immunoprecipitated AMPAR complexes isolated using an anti-GluA1 C-terminal antibody (A), an anti-phospho-GluA1 Ser831 antibody (P), or a secondary anti-GluA1 immunoprecipitation performed on the phospho-GluA1 flow-through fraction (PA). Samples were probed with antibodies against total GluA1, phospho-GluA1 Ser831, and PRRT1. Right, schematic workflow of the sequential immunoprecipitation strategy.

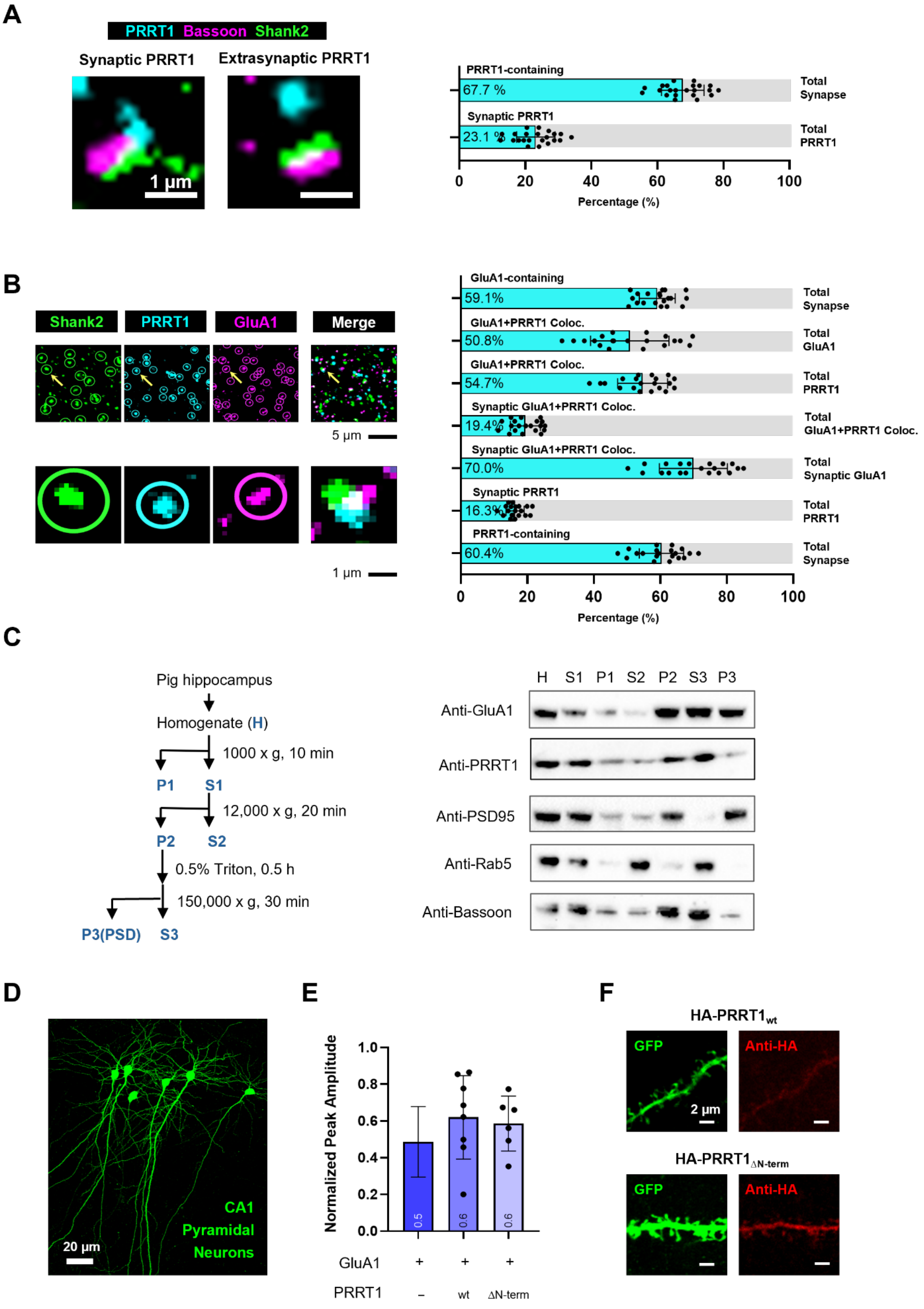

A
PRRT1 Bassoon Shank2
Synaptic PRRT1
Extrasynaptic PRRT1
1 µm
PRRT1-containing
67.7 %
Total Synapse
Synaptic PRRT1
23.1 %
Total PRRT1
0 20 40 60 80 100
Percentage (%)
B
Shank2
PRRT1
GluA1
Merge
5 µm
1 µm
GluA1-containing
59.1%
Total Synapse
GluA1+PRRT1 Coloc.
50.8%
Total GluA1
GluA1+PRRT1 Coloc.
54.7%
Total PRRT1
Synaptic GluA1+PRRT1 Coloc.
19.4%
Total GluA1+PRRT1 Coloc.
Synaptic GluA1+PRRT1 Coloc.
70.0%
Total Synaptic GluA1
Synaptic PRRT1
16.3%
Total PRRT1
PRRT1-containing
60.4%
Total Synapse
0 20 40 60 80 100
Percentage (%)
C
Pig hippocampus
Homogenate (H)
1000 x g, 10 min
P1 S1
12,000 x g, 20 min
P2 S2
0.5% Triton, 0.5 h
150,000 x g, 30 min
P3(PSD) S3
H S1 P1 S2 P2 S3 P3
Anti-GluA1
Anti-PRRT1
Anti-PSD95
Anti-Rab5
Anti-Bassoon
D
CA1
Pyramidal
Neurons
20 µm
E
Normalized Peak Amplitude
1.0
0.8
0.6
0.4
0.2
0.0
0.5
0.6
0.6
GluA1 + + +
PRRT1 – wt ΔN-term
F
HA-PRRT1wt
GFP
Anti-HA
2 µm
HA-PRRT1ΔN-term
GFP
Anti-HA


631

**Fig. S7. Localisation and function of PRRT1 in neurons**

**A**, Quantification of PRRT1 in expanded mouse hippocampal slice. Left: Confocal images of synaptic and extrasynaptic PRRT1 in brain slices stained with presynaptic marker Bassoon (magenta), postsynaptic marker Shank2 (green), and PRRT1 (cyan). True synapses were identified by colocalization of pre- and postsynaptic markers. Scale bar: 1 μm (biological scale: approximately 125 nm). Right: Bar graphs showing quantitative analysis of PRRT1 colocalization with synaptic markers. Three biological replicates were performed (independent expansion and staining trials, each from different slices from different adult mice, n = 22 image stacks).

**B,** Quantification of PRRT1 and GluA1 colocalization in expanded mouse hippocampal slice. Left: Representative processed confocal images showing spots identification (circle) in each channel based on pre-defined parameters. Lower panels are zoom-in images corresponding to regions indicated in the upper panels (yellow arrow). Scale bar for upper panels: 5 μm (biological scale: approximately 625 nm); Scale bar for zoom-in panels: 1 μm (biological scale: approximately 125 nm). Right: Bar graphs showing quantitative analysis of PRRT1 and GluA1 colocalization with synaptic marker Shank2. Three biological replicates were performed (independent expansion and staining trials, each from different slices from different adult mice, n = 20 image stacks).

**C**, Left, schematic workflow showing sequential centrifugation steps to isolate synaptosome fraction (H, homogenate; S, supernatant; P, pellet; fraction P2 containing synaptosome proteins; fraction P3 containing PSD-associated proteins) Right, western blot analysis of fractionated samples using antibodies against GluA1, PRRT1, PSD95, Rab5, and Bassoon.

**D,** Confocal images of electroporated CA1 pyramidal neurons in mouse organotypic hippocampal slice with GFP marker. Scale bar: 20 μm.

**E,** Bar graph showing normalised EPSC amplitudes from dual synaptic recordings in organotypic mouse hippocampal slice under baseline condition (GluA1, data included in Fig. 4C; GluA1+PRRT1$_{wt}$, n = 8; GluA1+ PRRT1$_{\Delta N\text{-}term}$, n = 6). Bars represent the geometric mean, and error bars indicate geometric SD.

**F,** Confocal images of transfected CA1 pyramidal neurons in mouse organotypic hippocampal slice. C-terminally HA-tagged PRRT1 wt and ΔN-term were electroporated with GFP as marker (green). Surface PRRT1 was visualized by immunostaining (red). Scale bar: 2 μm.

**Table S1. Cryo EM data processing and refinement statistics.**

| **Native GluA2 AMPAR** | A1A2A1A2 NTD (EMDB-58413) (PDB 31HF) | A1A2A3A2 NTD (EMDB-58414) (PDB 31HG) | A3A2A3A2 NTD (EMDB-58415) (PDB 31HH) | A2-AMPAR TMD (EMDB-58416) (PDB 31HI) | nonamer TMD (EMDB-58417) (PDB 31HJ) | decamer TMD (EMDB-58418) (PDB 31HK) | decamer NTD (EMDB-58419) |
|---|---|---|---|---|---|---|---|
| **Data collection and processing** | | | | | | | |
| Microscope | TFS Titan Krios | TFS Titan Krios | TFS Titan Krios | TFS Titan Krios | TFS Titan Krios | TFS Titan Krios | TFS Titan Krios |
| Detector | BioQuantum K3+GIF | BioQuantum K3+GIF | BioQuantum K3+GIF | BioQuantum K3+GIF | BioQuantum K3+GIF | BioQuantum K3+GIF | BioQuantum K3+GIF |
| Magnification | 105,000x | 105,000x | 105,000x | 105,000x | 105,000x | 105,000x | 105,000x |
| Voltage (kV) | 300 | 300 | 300 | 300 | 300 | 300 | 300 |
| Electron exposure ($e^-/Å^2$) | 42.8 | 42.8 | 42.8 | 42.8 | 42.8 | 42.8 | 42.8 |
| Defocus range (μm) | -1.2 to -2.4 | -1.2 to -2.4 | -1.2 to -2.4 | -1.2 to -2.4 | -1.2 to -2.4 | -1.2 to -2.4 | -1.2 to -2.4 |
| Pixel size (Å) | 0.826 | 0.826 | 0.826 | 0.826 | 0.826 | 0.826 | 0.826 |
| Symmetry imposed | C1 | C1 | C1 | C1 | C1 | C1 | C1 |
| Initial particle images (no.) | 758,375 | 758,375 | 758,375 | 758,375 | 758,375 | 722,448 | 722,448 |
| Final particle images (no.) | 445,187 | 216,321 | 96,867 | 758,375 | 65,517 | 48,206 | 48,206 |
| Map resolution (Å) | 3.1 | 3.1 | 3.4 | 2.7 | 3.1 | 2.9 | 4.0 |
| FSC threshold | 0.143 | 0.143 | 0.143 | 0.143 | 0.143 | 0.143 | 0.143 |
| Map resolution range (Å) | 2.6-7.0 | 2.7-8.0 | 2.8-8.5 | 2.4-6.5 | 2.6-7.0 | 2.5-7.5 | 3-7.2 |
| **Refinement** | | | | | | | |
| Initial model used (PDB code) | 7OCC | 7OCC, 9HPE | 7OCC, 9HPE | AF-predicted | AF-predicted | AF-predicted | - |
| FSC threshold | 0.143 | 0.143 | 0.143 | 0.143 | 0.143 | 0.143 | 0.143 |
| Map sharpening *B* factor ($Å^2$) | -107.2 | -138.2 | -86.3 | -76.6 | -52.5 | -43.1 | -75.3 |
| Model composition | | | | | | | |
| Non-hydrogen atoms | 13450 | 12967 | 11944 | 18990 | 18861 | 19742 | - |
| Protein residues | 1938 | 11715 | 1492 | 2403 | 2391 | 2502 | - |
| Ligands | 20 | 17 | 14 | 2 | 2 | 4 | - |
| *B* factors ($Å^2$) | | | | | | | |
| Protein (mean) | 102.25 | 97.49 | 127.87 | 150.57 | 56.84 | 169.81 | - |
| Ligand (mean) | 109.70 | 124.28 | 161.35 | 65.46 | 53.09 | 102.93 | - |
| R.m.s. deviations | | | | | | | |
| Bond lengths (Å) | 0.003 | 0.003 | 0.003 | 0.003 | 0.005 | 0.003 | - |
| Bond angles (°) | 0.567 | 0.556 | 0.659 | 0.639 | 0.695 | 0.669 | - |
| Validation | | | | | | | |
| MolProbity score | 1.33 | 1.40 | 1.59 | 2.08 | 1,79 | 1.77 | - |
| Clashscore | 4.86 | 4.63 | 7.93 | 8.83 | 10.00 | 8.78 | - |
| Poor rotamers (%) | 0 | 0 | 0 | 2.96 | 0 | 1.19 | - |
| Ramachandran plot | | | | | | | |
| Favored (%) | 97.65 | 97.05 | 97.15 | 96.29 | 96.12 | 95.68 | - |
| Allowed (%) | 2.35 | 2.95 | 2.85 | 3.66 | 3.88 | 4.32 | - |
| Disallowed (%) | 0 | 0 | 0 | 0 | 0 | 0 | - |

**Table S2. Outside-out current parameters measured for recombinant AMPARs. Values are mean ± s.d.**

| AMPAR constructs | $t_{deactivation}$ (ms) (n) | $t_{desensitisation}$ (ms) (n) | steady state (%peak) (n) |
|---|---|---|---|
| **GluA1** | 0.7 ± 0.1 (8) | 2.9 ± 0.5 (9) | 0.7 ± 0.3 (9) |
| **GluA1 + PRRT1** | 1.0 ± 0.3 (8) | 3.4 ± 0.6 (8) | 0.9 ± 0.5 (8) |
| **GluA1 + PRRT1$_{\Delta N\text{-term}}$** | 0.9 ± 0.2 (5) | 2.4 ± 0.4 (5) | 0.6 ± 0.5 (5) |
| **GluA1 + CNIH2** | 5.2 ± 1.4 (26) | 9.0 ± 2.2 (14) | 4.8 ± 1.9 (14) |
| **GluA1 + CNIH2 + PRRT1** | 6.1 ± 2.1 (19) | 10.0 ± 2.6 (11) | 6.1 ± 3.3 (11) |
| **GluA1 + CNIH2 + PRRT1$_{\Delta N\text{-term}}$** | 8.6 ± 2.5 (32) | 12.3 ± 3.5 (20) | 6.4 ± 2.5 (20) |
| **GluA1$_{C825S}$ + CNIH2** | 4.6 ± 1.8 (17) | 8.2 ± 2.3 (11) | 4.5 ± 1.9 (11) |
| **GluA1$_{C825S}$ + CNIH2+ PRRT1$_{\Delta N\text{-term}}$** | 5.1 ± 1.8 (18) | 8.7 ± 2.2 (14) | 4.0 ± 1.8 (14) |
| **GluA1$_{A3\ C\text{-tail}}$ + CNIH2+ PRRT1$_{\Delta N\text{-term}}$** | 6.2 ± 2.3 (13) | 11.9 ± 4.1 (7) | 7.0 ± 1.5 (7) |
| **GluA1 + γ8** | 2.5 ± 0.4 (9) | 7.5 ± 2.0 (9) | 3.2 ± 1.8 (9) |
| **GluA1 + γ8 + PRRT1** | 2.5 ± 0.74 (8) | 6.6 ± 0.9 (9) | 2.6 ± 1.7 (9) |
| **GluA1 + γ8 + PRRT1$_{\Delta N\text{-term}}$** | 2.5 ± 0.4 (7) | 6.2 ± 0.9 (7) | 1.8 ± 1.0 (7) |
| **GluA3 + CNIH2** | 5.4 ± 2.4 (10) | 26.0 ± 9.0 (10) | 11.6 ± 4.3 (10) |
| **GluA3 + CNIH2 + PRRt1** | 6.8 ± 3.1 (12) | 32.7 ± 8.8 (12) | 15.2 ± 7.9 (12) |
| **GluA3 + CNIH2 + PRRT1$_{\Delta N\text{-term}}$** | 6.2 ± 2.3 (14) | 30.4 ± 8.3 (14) | 13.9 ± 5.2 (14) |
| **GluA3$_{A1\ C\text{-tail}}$ + CNIH2 + PRRT1$_{\Delta N\text{-term}}$** | 9.5 ± 2.9 (12) | 30.1 ± 6.3 (7) | 17.3 ± 5.8 (7) |
| **GluA1/GluA2(R)_γ8 + CNIH2** | 4.1 ± 1.6 (6) | 16.7 ± 3.5 (6) | 7.6 ± 1.1 (6) |
| **GluA1/GluA2 (R)_γ8 + CNIH2 + PRRT1** | 4.1 ± 1.0 (6) | 19.7 ± 8.9 (6) | 9.1 ± 4.2 (6) |

**Movie S1.**

Supplementary Video 1. A rotating decameric structure of AMPAR complex. GluA1 (A/C positions) and GluA2 (B/D positions) are coloured blue and red, respectively. TARP-γ8 subunits at the B'/D' positions are shown in light green, CNIH2 subunits at the A'/C' positions in orange, and PRRT1 molecules in cyan. The zoomed-in view highlights the interaction between the resolved segment of the GluA1 C-terminal tail and the TM1 helix of PRRT1, including the C16 palmitoyl modification at C825.

**Data S1. (separate file)**

Raw data from electrophysiology recordings.
